# A Cortical Microcircuit for Region-Specific Credit Assignment in Reinforcement Learning

**DOI:** 10.1101/2024.10.15.618337

**Authors:** Quentin Chevy, Zoltan Szadai, Loreen Hertäg, Morgan Moll, Elizabeth T Gibson, Rui Ponte Costa, Balazs Rozsa, Adam Kepecs

## Abstract

The distributed architecture of the cortex poses a fundamental challenge for reinforcement learning: how to assign credit specifically to regions that contribute to successful behavior? Cortical neurons can be driven by both global reinforcers, like rewards, and local sensory features, making it difficult to disentangle these influences. To address this, we investigated cortical reinforcement learning by manipulating the reward-predictive sensory modality during learning tasks, while monitoring key regulators of cortical activity—local inhibitory neurons, and cholinergic inputs. We found that VIP interneurons are broadly recruited by reward-predictive cues via a modality-independent cholinergic signal. However, when task demands aligned with local computation, SST interneurons suppressed VIP recruitment through an inhibitory feedback loop. A computational model demonstrates that this cholinergic-VIP-SST interneuron circuit motif enables targeted reinforcement learning and region-specific credit assignment in the cortex. These results offer a neurobiologically-grounded framework for how the cortex uses global reinforcement signals to direct plasticity to task-relevant regions, enabling those regions to adapt and fine-tune their responses.

## Introduction

Reinforcement learning theory powerfully explains how animals —and artificial agents— adapt behavior to maximize rewards from their environment^1–3^. Neuronal correlates of reinforcement learning have been explored mainly in the basal ganglia, where dopamine neurons implement a reinforcement prediction error algorithm^4–7^. There, dopaminergic signals modulate striatal neuron activity and plasticity to support action selection guided by reinforcement feedback^7–10^. Although work on cortical plasticity is usually limited to unsupervised learning contexts, evidence of reinforcement feedback impacting neuronal computation throughout the cortical hierarchy is emerging^11^. However, the distributed and modular architecture of the cortex presents a challenge to any reinforcement learning algorithm: how to assign credit to specific regions of the cortex that contribute to successful behavior?

This challenge, known as the credit assignment problem, involves determining which cortical regions should be credited with influencing a behavior that led to a reward, and subsequently directing plasticity to those regions. Local inhibitory interneurons, which are integral to controlling the flow of information within cortical circuits, are ideally positioned to play a key role in this process because they regulate the activity of principal cells. Among these cortical inhibitory interneurons, vasoactive intestinal polypeptide-expressing (VIP) interneurons (VIP-INs) intriguingly show direct recruitment by primary reinforcers across the cortex, including in primary sensory areas^12–16^ — a phenomenon that challenges the conventional understanding of their role in local circuit function and raises questions about their broader involvement in cortical processing and reinforcement learning.

VIP interneurons are a subclass of inhibitory neurons located in the superficial layers of the cortex^18–20^. Rather than directly influencing pyramidal neurons, VIP-INs preferentially target other interneurons, providing a disinhibitory drive, boosting pyramidal neurons activity^13,21–24^. Despite their involvement in local sensory processing, VIP-INs are often characterized by weak tuning^13,16,25–30^, suggesting that their primary function might lie elsewhere. In contrast to their subtle role in sensory tuning, VIP-IN activity is strongly modulated by arousal and locomotion, positioning them as key mediators of top-down dependent plasticity and learning throughout cortex^31–40^. However, the source and impact of the reinforcement-related recruitment of VIP-INs remains poorly documented^15^. Whether VIP-INs merely signal the presence of a behavioral reinforcer or convey a more nuanced reinforcement-related signal remains an open question^12,14,17^.

VIP-IN recruitment by top-down feedback—arousal changes or primary reinforcers—is likely inherited from neuromodulator inputs to cortex. In particular, VIP-IN expression of nicotinic and muscarinic receptors allows for cholinergic-dependent modulation of VIP-IN activity both in vitro and in vivo^15,21,41–47^. Basal forebrain cholinergic projections orchestrate widespread top-down control of cortical state and processing^15,48–52^. Basal forebrain cholinergic neurons are not only implicated in arousal, movement, attention, and even time-related modulation of cortical activity^49,53–56^, but also encode primary and secondary reinforcers^15,52,57–62^. How these reinforcement prediction signals could support reinforcement learning, through VIP-IN recruitment in the cortex, remains largely unknown ^52^.

In this study, we set out to unravel the enigmatic role of VIP-INs in integrating global reinforcement feedback with local cortical processing. We used fiber photometry and 2-photon acousto-optic deflector (2P-AOD) microscopy to measure changes in interneuron activity and acetylcholine throughout the cortex during learning of tasks designed to separate local activity from reinforcement signals. We found that acetylcholine provides a global, sensory-independent reward expectation signal to the cortex. This global signal recruits VIP interneurons, leading to the inhibition of a subgroup of somatostatin (SST) interneurons during learning of reward predicting cues. We also identified a second group of SST interneurons likely responsible for turning off VIP interneuron disinhibition upon learning of locally-encoded reward predicting cues. Finally, we incorporated these findings into a model to show how disinhibition and feedback inhibition gate reinforcement-dependent learning of local inputs. These results lead us to propose that the acetylcholine-VIP-SST triad constitutes a canonical cortical motif, supporting credit assignment and reinforcement learning throughout the cortex.

## Results

### Reward expectation modulates both global and local recruitment of VIP interneurons

Cortical VIP interneurons (VIP-INs) are recruited by both local (*e.g.* auditory tuning in auditory cortex) and global signals (*e.g.* reward-mediated signals observed throughout cortex). Here, we asked how these local and global modes of recruitment are impacted by reward expectation. We used fiber photometry to measure the changes in fluorescence from GCaMP-expressing VIP-INs in the auditory cortex (**Figure 1c**) of head-fixed mice learning an auditory Pavlovian task (**Figure 1a-b**). The task consisted of learning the association between an auditory chirp (conditional stimulus, CS) and the delayed delivery of a reward (cued reward, CR). Throughout training, the animals were also presented with uncued water reward (UR) or neutral auditory stimulus (NS) that was never followed by a reward (see Methods). Learning of the task was determined by the development of stronger anticipatory licking to the CS than the NS (**Figure 1b** late CS vs. NS lick rate p<0.001, WSRT). Before training started, we measured the sensory (**Figure 1g and S1b**) and reward-evoked (**Figure S1c**) VIP-IN responses and determined that they were not significantly contaminated by hemodynamic artifacts (**Figure S1d-e-f,** p=0.12 and p=0.06 for sensory and reward respectively).

**Figure 1.**
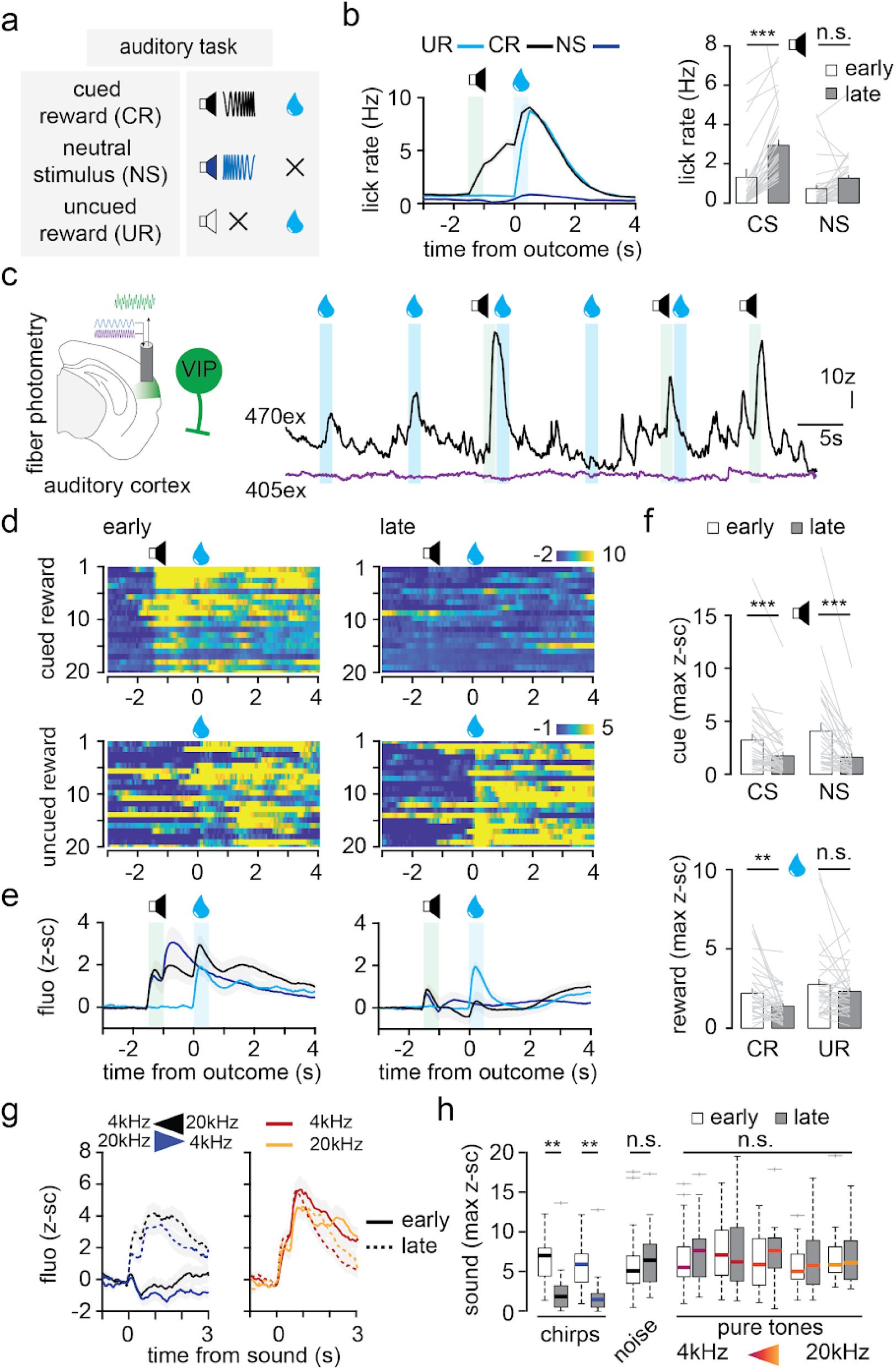
Reward expectation modulates sensory and outcome responses of auditory VIP interneurons. (a) Associative auditory Pavlovian task schedule. (b) Average lick rate (left) and quantification of anticipatory licking early and late in learning (right, n=34). (c) Diagram (left) and example trace of VIP-IN fiber photometry recordings in the auditory cortex during the task described in A (right). (d) Pseudo-colored raster plot showing z-scored VIP-IN single trial activity early (left) and late (right) in training. (e) Average VIP-IN activity early (left) and late (right) in training for trial types, colored as in B (n=34). (f) Change in average VIP-IN responses to cue (top) and reward (bottom). (g) Average VIP-IN activity mediated by chirps (left) or pure tones (right) early and late in training (n=22). (h) Change in average VIP-IN responses to sounds. CR: Cued Reward, CS: Conditional Stimulus, NS: Neutral Stimulus, UR: Uncued Reward. n: mice, **: p<0.01, ***: p<0.005, n.s.: non-significant.

VIP-INs in the auditory cortex showed a strong reduction of both sensory and reward responses during learning of the auditory Pavlovian task. On the first day of training, VIP-INs were recruited by both CR and UR (**Figure 1d-e-f**). Similarly, both CS and NS elicited comparable sensory responses of VIP-INs (**Figure 1f-g and S3)**. However, after learning, the recruitment of VIP-INs to CR decreased (**Figure 1f**, 63% of early peak fluorescence, p=0.009 WSRT), while the UR responses remained stable (p=0.2 WSRT). Unexpectedly, we also observed a strong decrease in the cue-evoked recruitment of VIP-INs for both CS and NS (**Figure 1f** 54%, p<0.001 and 40%, p<0.001 of CS and NS early peak fluorescence). Importantly, this decrease in sensory response was not observed for other sound types such as pure tones or white noise which were never presented during training (**Figure 1g-h** p=0.815 ranked ANOVA). The decrease in response to both CS and NS, but not to other sounds, could either reflect stimulus generalization, or how the auditory cortex participates in discriminating between CS and NS during this task.

Associative training using complex sounds is known to differentially recruit cortex as compared to using pure tones^63^ but we did observe similar results using pure tones predicting reward delivery (**Figure S3,** 42% of early CS peak fluorescence p<0.05).

The decreased response to predicted reward suggests that VIP-INs encode reward *expectation* rather than simply indicate the presence or the absence of reward. However, the decrease in *cue*-evoked recruitment of VIP-INs during learning suggests a more complex coding of cue-predicting reward.

### VIP interneurons encode reward-predictive cues in absence of local sensory drive

Removing local sources of VIP-IN recruitment revealed a conditional cue coding by VIP-INs in the auditory cortex. To better understand the reward-predictive code of auditory VIP-INs, we sought to eliminate the influence of confounding local auditory sensory inputs while still providing an outcome-predictive cue. Thus, we switched to a visual-based associative task while recording GCaMP-expressing VIP-INs in the auditory cortex using fiber photometry (**Figure 2a-b and figure S4**, late CS vs. NS lick rate p<0.001, WSRT).

**Figure 2.**
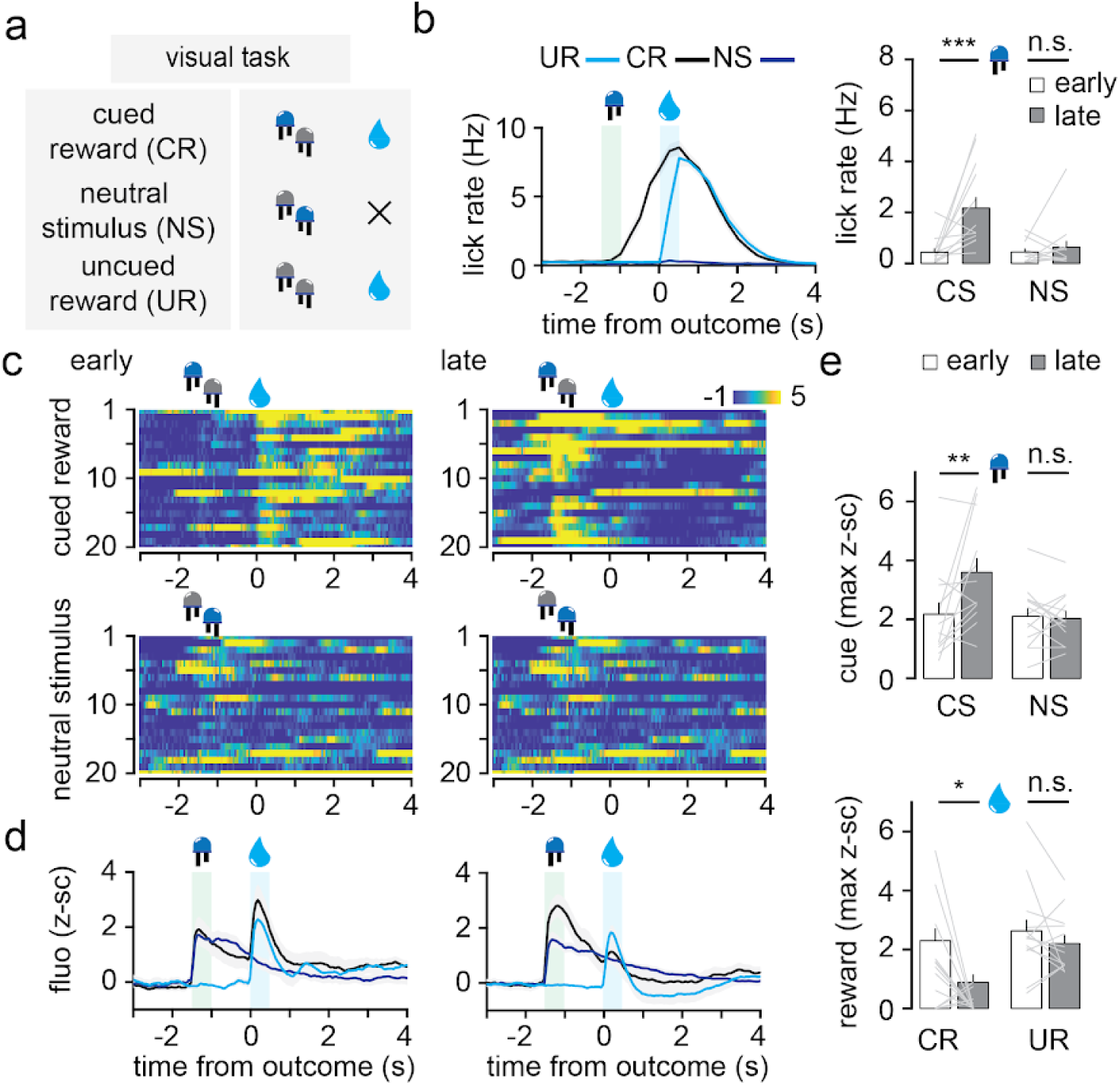
VIP interneurons encode reward predicting visual cues in auditory cortex. (a) Associative visual Pavlovian task schedule. (b) Single animal average lick rate after learning (left) and quantification of anticipatory licking early and late in learning (right, n=13). (c) Pseudo-colored raster plot showing z-scored VIP-IN single trial activity early (left) and late (right) in training. (d) Average VIP-IN activity early (left) and late (right) in training for trial types, colored as in B (n=13). (e) Change in average VIP-IN responses to cue (top) and reward (bottom). CR: Cued Reward, CS: Conditional Stimulus, NS: Neutral Stimulus, UR: Uncued Reward. n: mice, *: p<0.05, ***: p<0.005, n.s.: non-significant.

During learning of the visual task, we found that the expectation-dependent recruitment of VIP-INs during reward delivery was identical to that observed for the auditory task (**Figure 2c-d-e**): CR responses decreased with learning (38% of early peak fluorescence, p=0.02), while the UR responses remained stable (p=0.3). However, the dynamics of the cue-evoked responses were different between the auditory and the visual tasks. On day 1, VIP-INs showed a small recruitment by both visual stimuli (**Figure 2d-e**, CS vs. NS, p=0.7). We hypothesized that these responses could represent a saliency-dependent recruitment (see also **Figure 5 and 7** and Discussion). After learning, VIP-INs showed an increase in recruitment specifically to the CS (165% of early peak fluorescence, p=0.02), while the NS-evoked responses did not change (p=0.8). Manipulating the sensory modality of the reward-predictive cues allowed us to separate learning-dependent local and global recruitment of VIP-INs in the auditory cortex. We then pursued our investigation of reward prediction coding by VIP-INs by asking whether it could be observed in other cortical areas.

### Reward predicting cue coding is limited to reward-responsive VIP interneurons

While showing that reward predicting cue coding by VIP-INs is generalizable to other cortical areas, we also found that this coding is specific to reward-responsive VIP-INs. We recorded VIP-INs in the medial parietal cortex (mPTA), where we previously reported that reward- and punishment-mediated VIP-IN recruitment^16^. We used 2-photon acousto-optic deflector (2P-AOD) microscopy^16,64,65^ to access single VIP-IN activity throughout the cortical layer of the mPTA in animals trained on the auditory task (**Figure 3a**). Similarly to what we previously reported, we found that most mPTA VIP-INs responded to reward (84% of recorded VIP-INs, **Figure 3b**). We separately analyzed the reward-responding (VIPp) and non responding (VIPn) cells after learning of the auditory task (**Figure 3c-d**). We found that VIPp-INs had stronger responses to CS than NS (p<10^-06, KS). VIPn-INs (those that did not respond to reward) had smaller responses to CS than VIPp-INs (p<10^-08, KS, **Figure 3e**). Just as we observed in the auditory cortex, VIP-INs were less recruited by cued than uncued reward (p=0.006, KS).

**Figure 3.**
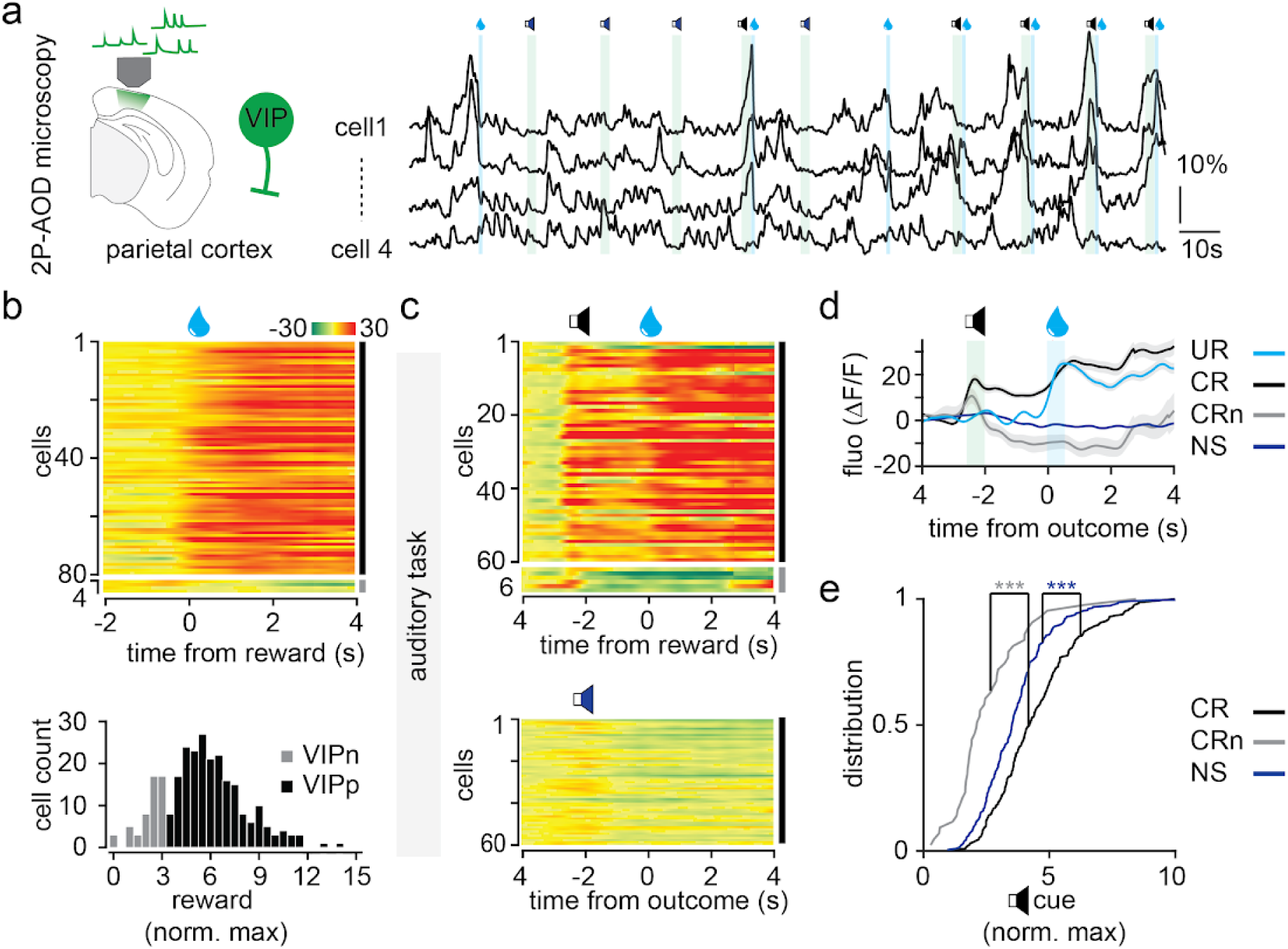
Reward-responsive VIP interneurons encode reward predicting auditory cues in parietal cortex. (a) Diagram (left) and concatenated trace of single VIP-IN AOD 2-photon microscopy recordings in the parietal cortex (right). (b) Pseudo-colored raster plot showing average z-scored fluorescence from single VIP-IN during uncued reward trials (top). Distribution of the average VIP-IN reward responses used to separate reward responsive (VIPp) and non-responsive (VIPn) neurons (bottom) (c) Pseudo-colored raster plot showing average z-scored fluorescence from single VIP-IN during CR and NS trials. VIP-IN were separated in VIPp and VIPn as in b. (d) Average VIP-IN activity for the data shown in c. Data for UR, CR and NS are from VIPp subpopulation, CRn shows the average cued reward activity of VIPn. (e) Single VIP-INs average responses to cues, pseudo-colored as in d (n=227/43 VIPp/VIPn). CR: Cued Reward, NS: Neutral Stimulus, UR: Uncued Reward. n: mice, ***: p<0.005.

Reanalyzing our previous work that characterized VIP-IN reward responses across many cortical regions also revealed learning-related reward and cue modulation of VIP-INs ^16^. Previously, we recorded single VIP-INs located in the dorsal cortex (PTA, motor, visual and somatosensory areas) using 2P-AOD microscopy during an auditory go-nogo task. Although these data were not longitudinal (*i.e.* not recorded across consecutive days), some animals’ performance improved between early and late trials within the same session (d’ early vs. late, 50+/-1% vs. 80+/-1%, p=0.004). Recordings from these animals showed that VIP-IN responses switched from the reward to the auditory cue delivery (reward/cue DF/F early 1.3+/-0.3 vs. late 0.7+/-0.1, p=0.02, **Figure S5c**). For the animals whose performance did not improve, VIP-IN activity did not change between the early and late trials in the training session (**Figure S5c**).

So far, we have shown that VIP-INs encode reward predicting cues in the absence of local cortical engagement, across all recorded cortical areas. However, when task demands are aligned with local cortical processing, VIP-IN recruitment by local inputs decreases after learning. These observations raise two critical questions: first, what mechanisms underlie the global ability of VIP-INs to encode predictions about rewards? Second, what accounts for the suppression of local VIP-IN activity when local neurons are engaged by the task’s sensory modality?

### Acetylcholine release signals reward delivery to VIP interneurons

We hypothesize that VIP-IN reinforcement responses are driven by basal forebrain cholinergic inputs to the cortex. Cholinergic basal forebrain neurons encode both primary reinforcers and reward prediction signals^52,59,62^, as well as recruit cortical VIP-INs^15,41^. Thus, we first asked whether cholinergic inputs to the cortex inform VIP-INs of the delivery of a reward.

We took advantage of the recombinant fluorescent muscarinic receptor GRAB-ACh4.3^66^ to measure the release of acetylcholine (ACh) in the vicinity of tdTomato-expressing VIP-INs in the parietal cortex using 2P-AOD microscopy (see Methods). At baseline, most cholinergic-related fluorescence could be observed in the upper layers of cortex, where the majority of VIP-INs are (**Figure 4a**). Reward delivery elicited a release of ACh nearby VIP-INs (**Figure 4b**). We replicated this observation in the auditory cortex using bilateral fiber photometry recordings and observed that ACh release increased upon reward delivery with similar dynamics to the reward-mediated recruitment of VIP-INs (**Figure 4c-d**, p=0.77 combining bilateral and independent GRAB-ACh, VIP-GCaMP recordings, see also **Figure S9a and S9b**). Reward-evoked ACh transients appeared to ride on top of a tonic ACh presence related to wakefulness (**Figure S6a-b**) and were not observed using the mutant ACh-insensitive version of GRAB-ACh (**Figure S6c**). Air puff punishment also led to a strong release of ACh in the auditory cortex (**Figure S6d**). Reward-evoked ACh release directly onto cre-expressing VIP-IN could also be detected using a cre-dependent GRAB-ACh virus (**Figure S6e**).

**Figure 4.**
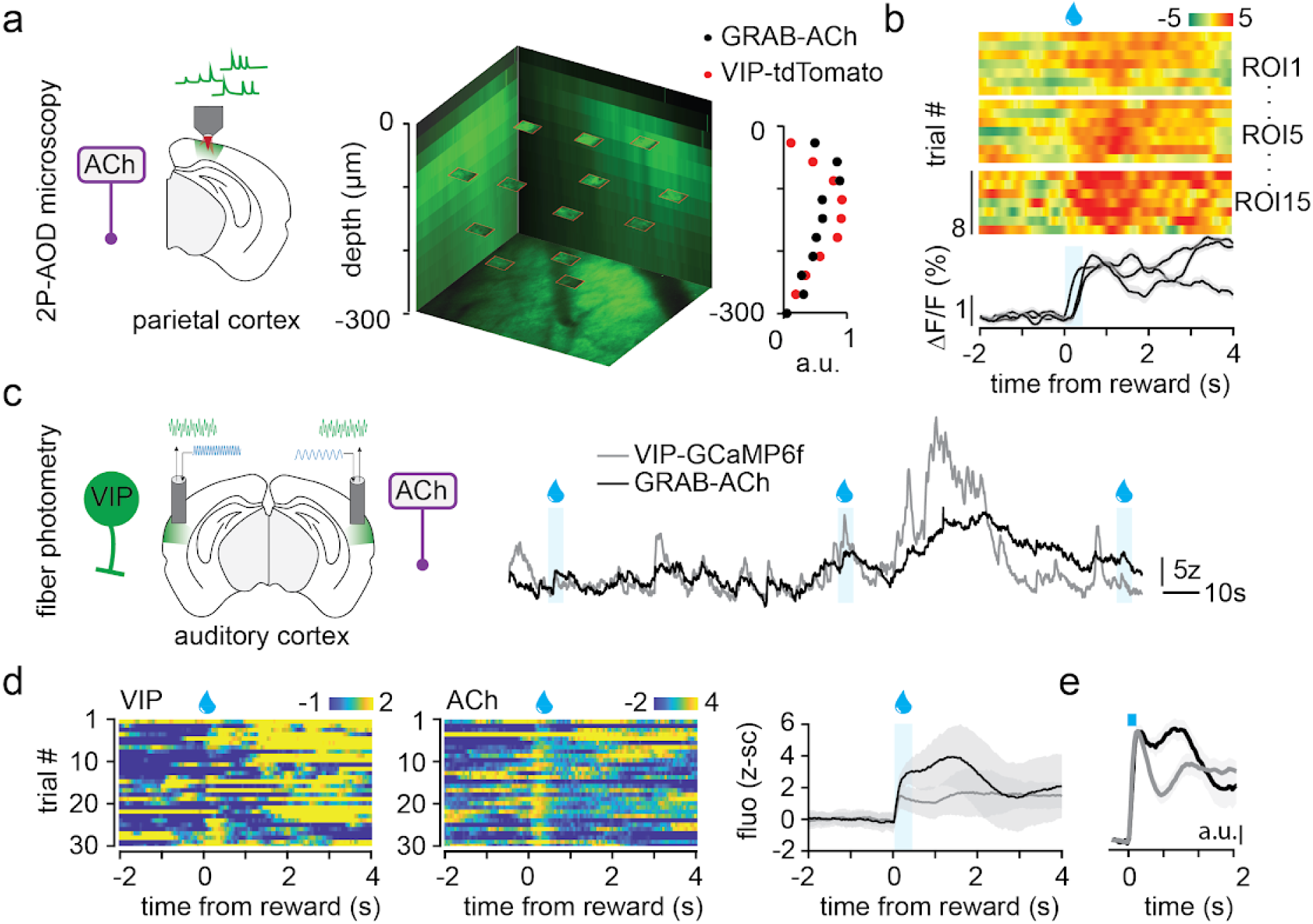
Acetylcholine release signals reward to cortical VIP interneurons. (a) Diagram (left) and 3D projection (middle) of AOD 2-photon microscopy recordings of acetylcholine release in the vicinity of tdTomato-expressing VIP-INs in parietal cortex. GRAB-ACh-related fluorescence and VIP-IN relative density (right). (b) Example raster plots (individual region of interest ROI) and mouse-average fluorescence traces showing the reward-evoked release of ACh (n=3). (c) Diagram (left) and example traces (right) from bilateral fiber photometry recordings of VIP-INs and ACh release in auditory cortex. (d) Pseudo-colored z-scored fluorescence raster plot and average reward-related activity for co-recorded VIP-IN and GRAB-ACh (n=11). (e) Normalized reward-evoked transient for GRAB-ACh (n=19) and VIP-IN recordings (n=55). Normalized slope was calculated over the blue square period. a.u.: arbitrary unit, n: mice.

Finally, we found that inhibiting cholinergic inputs to the cortex decreased the recruitment of VIP-INs by reward and punishment, similar to the functional relationship between cholinergic inputs and VIP-IN activity observed in motor cortex^15^. To acutely inactivate cholinergic inputs, we targeted a light-switchable inhibitory opsin SwitChR^67^ to cholinergic neurons in the basal forebrain, and injected a cre-dependent GCaMP6f virus in the motor cortex of a VIP-cre x ChaT-cre mouse line. We used 2P-AOD to monitor single VIP-IN responses to reinforcers (see task structure in **Figure S5a**) with or without inhibiting basal forebrain cholinergic neurons. Inhibiting cholinergic inputs to the motor cortex using SwitChR stimulation led to a 33.3% reduction in reinforcement-mediated activation of VIP-INs (**Figure S6e-f** p<0.001). The local cholinergic signals recorded with 2P-AOD microscopy were strikingly similar in both amplitude and dynamics to those recorded using fiber photometry in auditory cortex and mPFC (**Figure 4 and S8**). These similarities highlight the volumetric and broadcasting nature of the ACh reinforcement signal, which could in turn support the global recruitment of VIP-INs by primary reinforcers.

### Acetylcholine provides a global reward prediction coding to cortex

We found that acetylcholine release in cortex encodes reward predicting cues irrespective of the sensory modality. The recruitment of basal forebrain cholinergic neurons by reward is modulated by behavioral expectation^61,62^. However, it is mostly unknown whether this expectation modulation is also present in cortex^68^, or could be customized to targeted cortical areas or sensory modalities^49,51,61^.

To answer those two points, we monitored ACh in the auditory cortex (**Figure 5 and S7**) and the medial prefrontal cortex (mPFC, **Figure S8**) while mice were trained on either the auditory task (**Figure 5b**, CS licks early vs. late, p<0.005) or the visual task (**Figure 5b**, CS licks early vs. late, p<0.005).

Early in training, both CR and UR led to a similar release of ACh for both task modalities (p=0.24 and p=0.27 for auditory and visual task respectively). The cue presentations also led to a small release of ACh that was identical for both CS and NS (CS vs. NS: p=0.2 and p=0.4 for auditory and visual task respectively), reminiscent of the one observed for VIP-INs early in the visual task (**see Figure 2**). The similarities between the two tasks continued late in training: the release of ACh increased for both auditory and visual CS (**Figure 5i**, 670%, p<0.005 and 169%, p<0.005 of early average fluorescence, for auditory and visual task respectively), while the NS elicited the same amount of ACh release as on day 1 (p=0.43 and p=0.416 WSRT for auditory and visual task respectively). The ACh release during CR was strongly decreased after learning for both tasks (19%, p<0.005 and −6% p<0.05 of early average fluorescence for auditory and visual task respectively). Similar results were observed when looking at the residual fluorescence changes upon cue and reward delivery (**Figure 5e-h-j**, see also methods).

We observed the same reward prediction coding by ACh release in the auditory cortex using an olfactory-based task (as in Sturgill et al., 2020, **Figure S8c to f**). Recording ACh release in another cortical area such as mPFC during either the visual or the olfactory version of the associative task led to the same reward prediction phenomenology: increase in conditional cue-evoked ACh release and no additional release upon predicted reward delivery (**Figure S8g-h**).

Thus, ACh release in cortex appears to convey a global, sensory-modality independent reward prediction signal that could drive reward-mediated VIP-IN activity during learning. However, the global signal provided by cortical cholinergic inputs cannot explain the inhibition of auditory VIP-IN recruitment by local sensory inputs during learning of the auditory task.

### SST interneuron recruitment by reward is heterogeneous

VIP-INs both inhibit and are inhibited by somatostatin interneurons^21,24^ (SST-INs). While the former drives the classical plasticity gating function of VIP-INs, we hypothesized that the latter could support the learned suppression of VIP-IN local activity (**Figure 1**). Thus we decided to record SST-INs during learning of the visual and auditory tasks.

We first sought to characterize the sensory and reward responses of SST-INs, and although fiber photometry typically yields averaged signals, we found unexpected heterogeneity in SST-INs responses across animals. Recordings clustered into two types: 15 of 20 animals showed strong recruitment of auditory SST-INs by reward (we denote these SSTp-INs) while the remaining 5 animals showed minimal or no response to reward delivery (SSTn-INs, **Figure 6c**, p<0.001 MWRST). This heterogeneity could not be explained by a difference in recording quality, as both SSTp-INs and SSTn-INs showed similar auditory-evoked responses in the auditory cortex (**Figure 6d**, p=0.75). We hypothesize that the variability in reward-related activity among SST-INs may be due to differences in SST-IN subtypes across cortical layers. Variations in the placement of fiber photometry probes in the auditory cortex likely contributed to this variability, although histological analysis could not quantify it (not shown). However, we found evidence of this putative layer-dependent reward responses of SST-INs using 2P-AOD microscopy recordings in the primary visual cortex (**Figure 6f**). We found that deeper SST-INs responded more strongly to reward than superficial ones (Pearson correlation=0.38, p<0.005 **Figure S9g**). A clustering analysis further identified three distinct SST-IN subgroups based on their reward responses: inhibited (SSTn-IN), fast (f-SSTp) and slow responding (s-SSTp) interneurons (**Figure 6f**). SSTn-INs were preferentially located in the superficial layers of the visual cortex (depth_SSTn_=-163 µm, n=13 cells) when compared to f-SSTp-IN and s-SSTp-IN subgroups (depth_f-SSTp_=-310 µm, n=29 cells, depth_s-SSTp_=-343 µm, n=65 cells). We suggest that SSTn-INs could be preferentially targeted by superficial VIP-INs leading to their overall inhibition during reward delivery.

**Figure 5.**
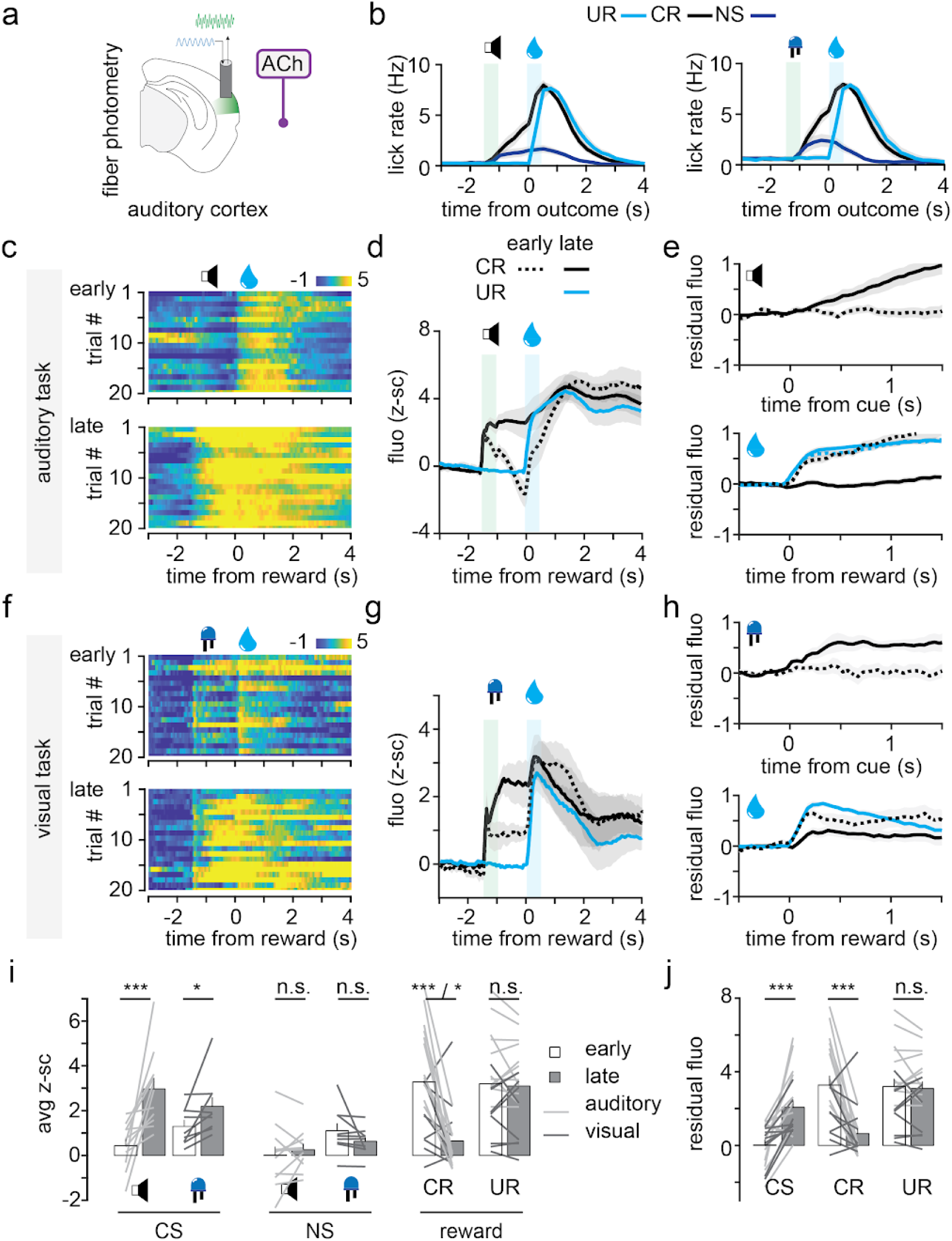
Sensory-modality independent reward prediction coding by acetylcholine in auditory cortex. (a) Diagram of fiber photometry recordings of ACh release in the auditory cortex. (b) Average lick rate after learning of the auditory (left, n=11) and visual task (right, n=9). (c) Pseudo-colored z-scored GRAB-ACh fluorescence raster plots for cued reward trials on the first (top) and last day (bottom) of training of the auditory task. (d) Average z-scored fluorescence changes during UR and CR (n=11). (e) Residual fluorescence (see Methods) during the cue (top) and reward (bottom) periods. (f-g-h) Same as (c-d-e) but for the visual task (n=9). (i) Changes in cue- and reward-related GRAB-ACh fluorescence for the two tasks. (j) Changes in average residual fluorescence during CS, CR and UR. Data from both tasks were pulled and analyzed together. n: mice, n.s.: non-significant, *: p<0.05 **: p<0.01 ***: p<0.005.

**Figure 6.**
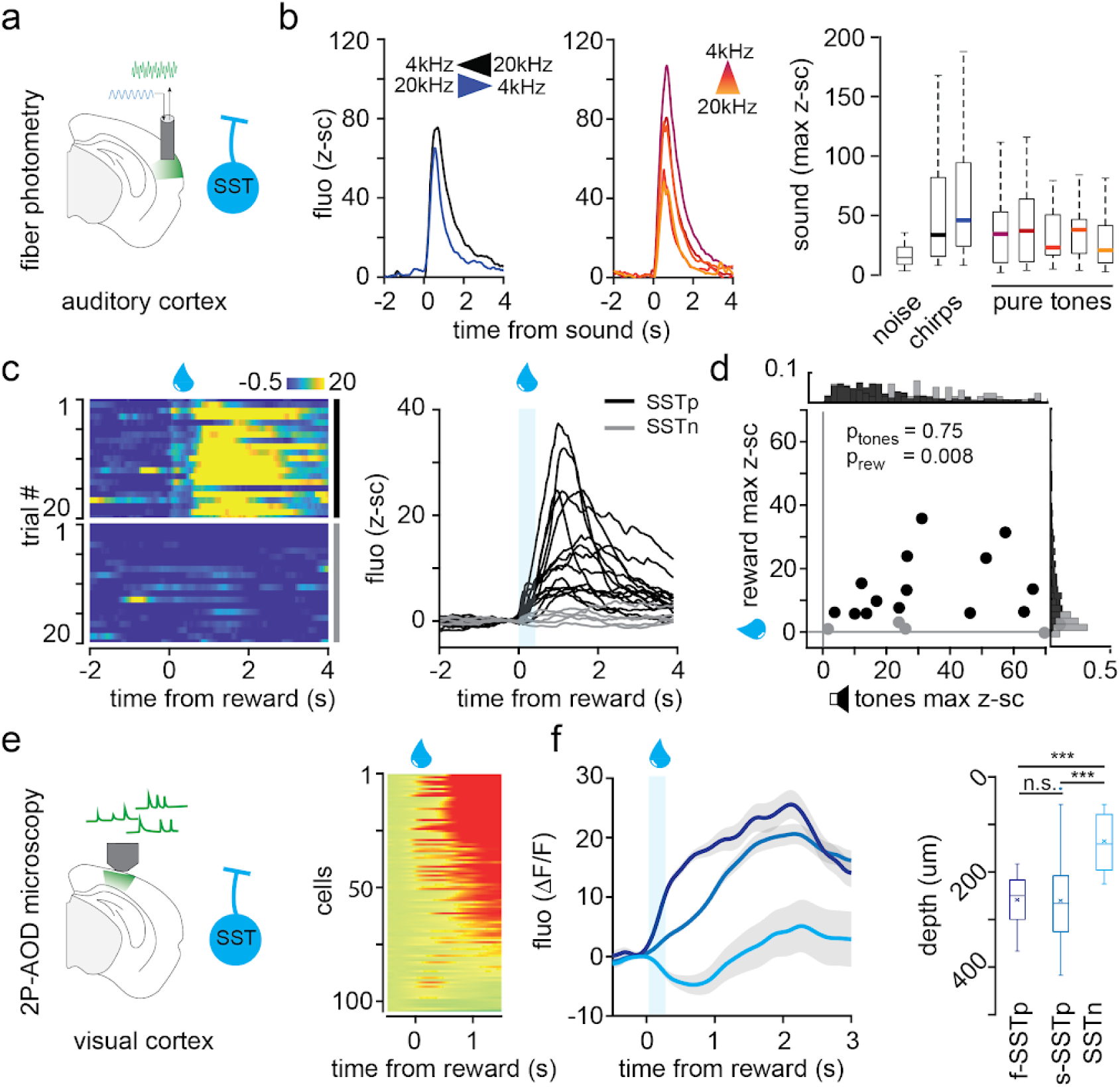
Cortical layer-related heterogeneity in SST interneuron reward responses. (a) Diagram of the fiber photometry recordings of SST-INs in the auditory cortex. (b) Average z-scored fluorescence traces and quantification of SST-IN sound-evoked activity (n=20). (c) Single trial (left) and average z-scored fluorescence recordings (right) of reward responsive (SSTp, black, n=15) and reward non-responsive (SSTn, gray, n=5) SST interneurons. (d) Comparison of the maximum reward- and pure tone-evoked responses of SST-INs. Outside bar plots show the distribution of individual responses. (e) Diagram (left) and average single cell reward responses of SST-INs (right) recorded using AOD 2-photon microscopy in visual cortex. (f) Clustering analysis separating SST-IN between slow (s-SSTp, n=65 cells) and fast (f-SSTp, n=29 cells) reward responding and reward non responding (SSTn, n=13 cells) (left). Correlation between cluster identity and depth (right). n: mice (b to d), cells (e to f). n.s.: non-significant, **: p<0.01 ***: p<0.005.

In addition to the diversity in reward response magnitude, we also observed that the population (**Figure S9a-b**) and single cell (**Figure S9d-e**) SST-IN reward responses were generally delayed when compared to those of VIP-INs and ACh (fiber photometry: p<0.001, F=13.9 ANOVA. 2P-AOD: SST vs. VIP p<0.001 MWRST). Importantly, this delay was not observed for auditory responses, ruling out variations due to GCaMP expression (**Figure S9b,** p<0.01). The delayed responses of SST-INs are unlikely to be driven by ACh inputs directly onto SST-INs, as ACh showed faster dynamics (**Figure S9g-h**, p<0.005).

Although the origin of these delayed responses is unclear, some theoretical studies make SST-INs the entry point for hierarchical reinforcement learning through back propagation^69^. In this model, SST-INs in higher cortical areas should respond faster to reward than SST-INs in primary sensory areas. We tested this hypothesis by recording SST-INs in mPFC, and observed that the SST-INs indeed showed a faster recruitment by reward in mPFC than in the visual cortex (**Figure S9g-h**, p<0.001). Another possibility is that the delayed recruitment could originate from (disinhibited) pyramidal cells locally recruiting the SSTp-IN subgroup. More experiments are necessary to narrow down the exact mechanism recruiting SST-INs during reward delivery.

### SST interneuron subtypes show opposite reward predicting cue coding

Next we considered whether the task-related activity of these two functional SST-IN types could explain the dynamics of VIP-INs during learning. We trained mice in either the visual or auditory task while recording auditory SST-INs using fiber photometry (**Figure 7**). We separated the animals based on the reward response of SST-IN (SSTp-IN vs. SSTn-IN, see above). Early in training, both groups showed similar recruitment to auditory cues (p=0.641 ranked ANOVA) and neither showed an increase in recruitment to the visual cues. Later in training however, the cue responses of the two SST-IN groups showed opposite dynamics during learning (**Figure 7c to f**). First, SSTp-INs increased their activity following the CS presentation for both auditory and visual cues (auditory CS n_early_=200 vs. n_late_=340 trials, p<0.005 - visual n_early_=225 vs. n_late_=225 trials, p<0.001 KS). Interestingly, SSTp-INs never directly responded to the visual cue presentation, but instead showed a ramping activity to the reward delivery. Second, SSTn-INs were less recruited by the auditory CS (n_early_=155 vs. n_late_=165 trials, p<0.001 KS), and showed a sustained inhibition following the visual CS presentation (n_early_=93 vs. n_late_=168 trials, p<0.05 KS).

### ACh-VIP-SST triad implements reinforcement-guided learning in cortex

All together, our results suggest the existence of an inhibitory feedback loop between VIP-INs and the two types of SST-INs that engages during reinforcement learning (**Figure 8a**). We developed a computational model to understand the role of this feedback loop in mediating cortical plasticity through the integration of global reinforcement feedback and local inputs. We simulated a rate-based network with an excitatory principal cell and three types of inhibitory neurons: SSTn, SSTp, and VIP (**Figure 8a**). In this model, the principal cell is tasked with aligning a local sensory input to a specified target directed to an apical-like dendritic compartment. At baseline, both SST-IN subtypes participate in canceling out this target input, thereby precluding learning at the principal cell. Driven by our experimental findings, we hypothesized that SST-IN subtypes are controlled differentially. First, the SSTn-IN is inhibited by the VIP-IN, as well as receives feedforward local input from it. Second, the SSTp-IN is recruited by the output of the principal cell and provides feedback inhibition onto both the VIP- and SSTn-IN. In our model, only the VIP-IN receives the reinforcement-carrying cholinergic input, although other cell types are likely to also benefit from cortical acetylcholine (see Discussion). Finally, the ACh dynamics during training were modeled as a simple sigmoid function to emulate the learning-related increase in cholinergic inputs during the reward-predictive cue period. We then tracked the cue response dynamics of the VIP-IN, SST-IN, and principal cell (**Figure 8b**).

**Figure 7.**
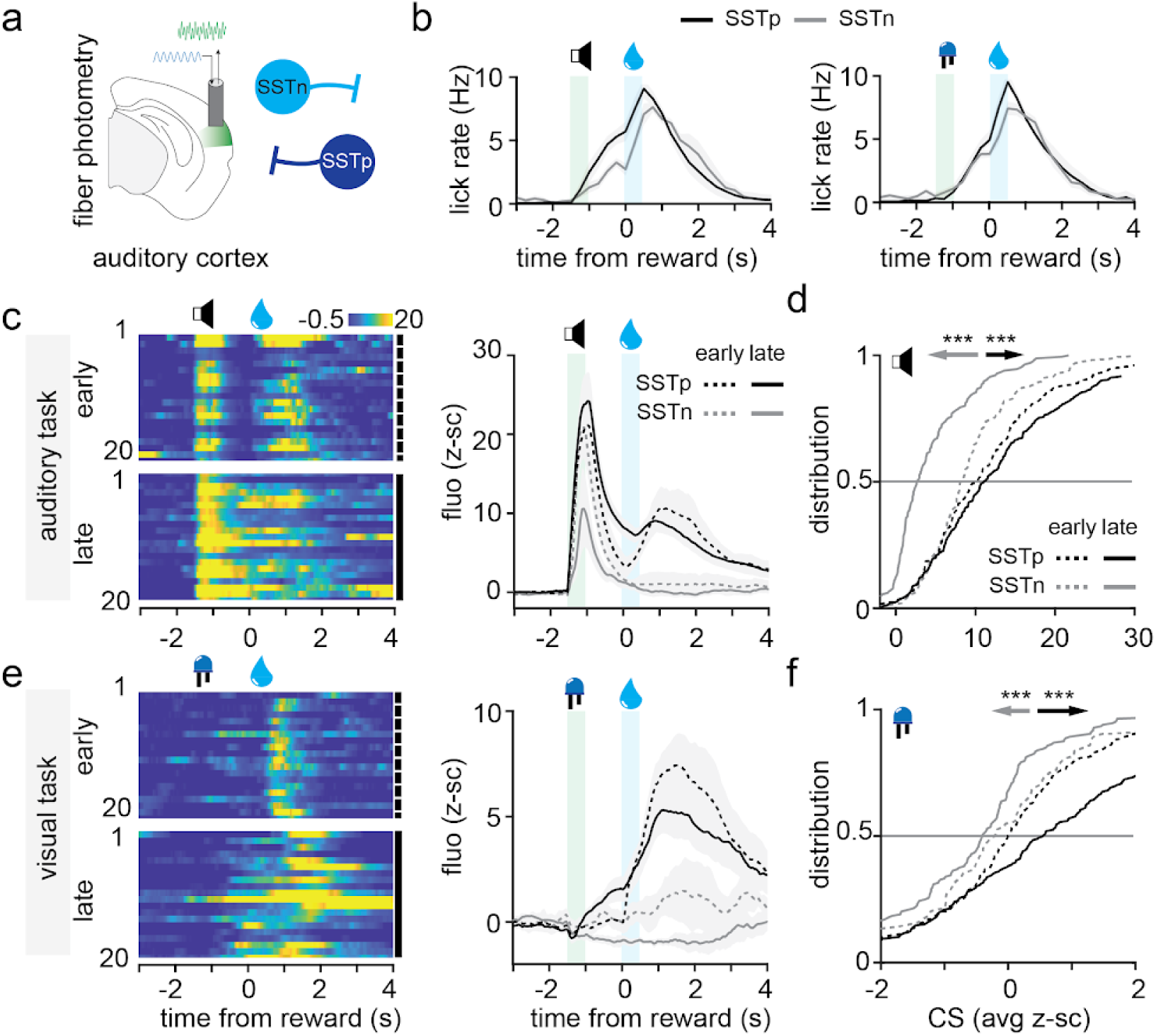
SST interneuron subtypes differentially encode reward predicting cues. (a) Diagram of the fiber photometry recordings of reward-responsive (SSTp-IN) and non-responsive (SSTn-IN) SST interneurons in the auditory cortex. (b) Average lick rate for cued reward trials late in learning of the auditory (top) and visual (bottom) tasks. (c) Pseudo-colored raster plot showing z-scored SSTp-IN single trial activity during auditory task training (left). Average z-scored fluorescence from SSTp-INs (n=12) and SSTn-INs (n=5) during cued reward trials early and late in training (right). (d) Changes in single trial CS responses early and late in training from SSTn-IN (155 vs.165 trials) and SSTp-INs (200 vs. 340 trials). (e-f) Same as (c-d) but for the visual task (e: n=9/n=3, f: 93 vs. 168 and 225 vs. 225 trials). n: mice, **: p<0.01 ***: p<0.005.

**Figure 8.**
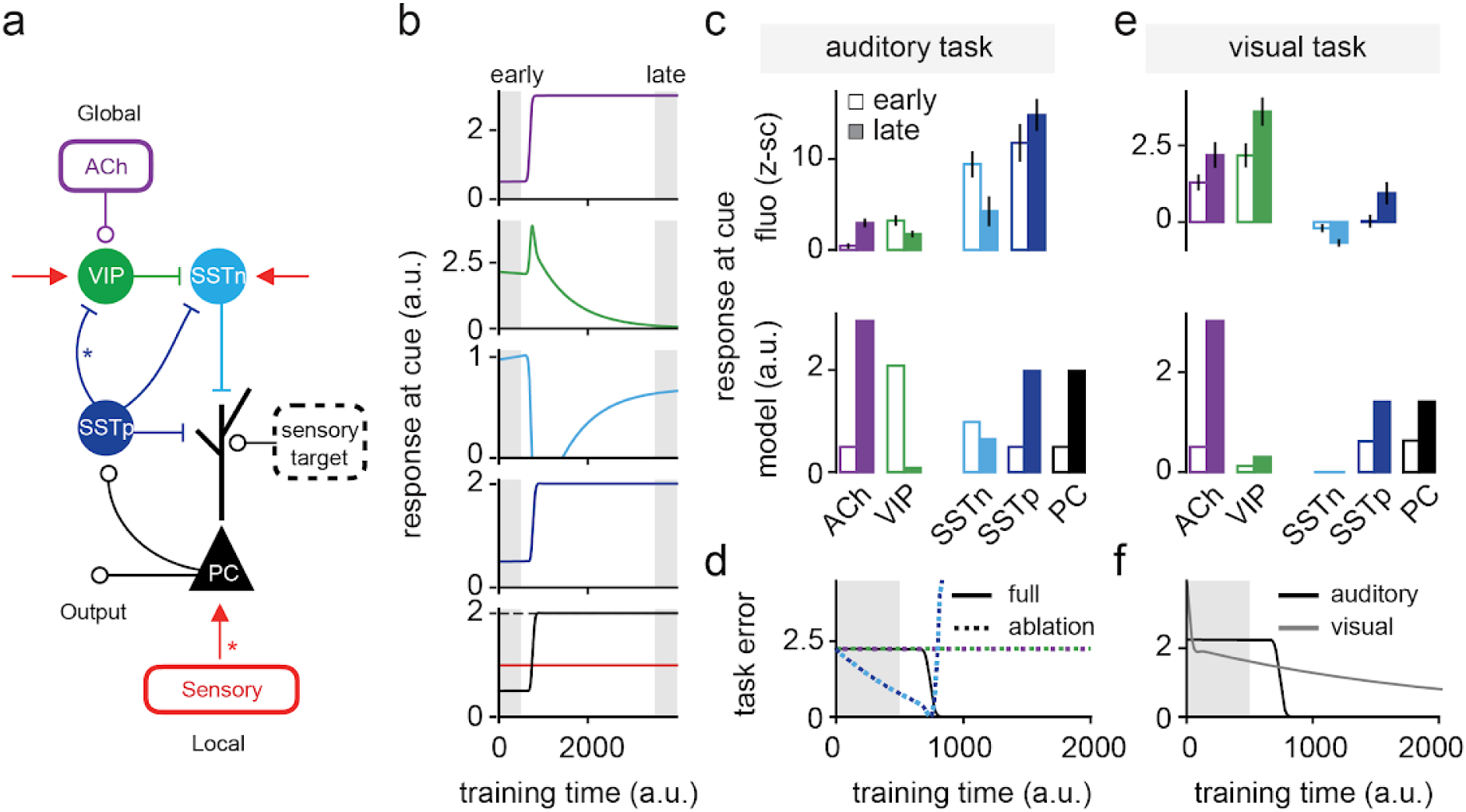
Control of VIP-SST microcircuit by global reinforcer enables task-specific plasticity. (a) Diagram of the computational model for reward-guided learning in cortical microcircuits. Red arrows indicate local, feedforward inputs, stars indicate plastic synapses. (b) Dynamics of ‘cue’ responses in the model for each neuronal cell-type as shown in a. (c) Comparison of the data and modeled cue responses from each neuronal cell type early and late in training for the auditory task. (d) Comparison of the full model performance with the task performance when acetylcholine (purple), VIP-IN (green), SSTp-IN (dark blue) or SSTn-IN (light blue) are ablated from the model. (e) Comparison of the data and modeled cue responses from each neuronal cell-type early and late in training for the visual task. (f) Comparison of the full model performance in the presence of strong or weak local inputs (e.g. auditory vs. visual task in auditory cortex).

During model training, we found that the increase in cholinergic input led to a disinhibition of the principal cell’s apical dendrite by recruiting the VIP-IN, thereby inhibiting the SSTn-IN (**Figure 8b**). This opened a window for learning, in which the dendrite encoded a prediction error, *i.e.*, the target minus the principal cell activity. Upon learning, the increase in the principal cell activity led to an increase in SSTp-IN activity. This, when combined with a local SSTp-to-VIP-IN plasticity rule (see Methods), fully inhibits VIP-IN activity. As a result, SSTn-IN are released from VIP-IN inhibition, reinstituting the inhibition of the apical dendrite, and thereby closing the plasticity window. These findings were robust across a range of local connectivity and learning rate parameters (**Figure S10 and S11**). Our model recapitulates our empirical data from VIP- and SST-INs in auditory cortex during the auditory associative learning task (**Figure 8c and Figure 1**): (1) decreased VIP-IN and SSTn-IN cue responses and (2) increased SSTp-IN cue response.

To demonstrate that the feedforward disinhibition and the feedback inhibition pathways respectively open and close a learning opportunity, we ablated individual cell types from our model and tested learning (**Figure 8d**). Removing key elements disrupted learning in distinct ways. Ablating the VIP-IN or acetylcholine inputs prevented plasticity altogether, eliminating the ability to learn cue-reward associations. Removing SSTp- or SSTn-INs caused overshooting of the target representation, because the dendritic inhibition of only one of the two SST-INs was not sufficient to balance the feedback target (**Figure 8d**). Finally, we tested how this model responds to small local inputs, akin to coding of visual associative learning task variables in the auditory cortex (**Figure 2**). We found that providing only reinforcement signals without meaningful sensory inputs dramatically reduced learning (**Figure 8f and S12**). This manipulation effectively decoupled ACh and VIP-IN activity from the rest of the network, revealing reinforcement coding from VIP-INs and recapitulating our data (**Figure 8e**). Overall, we demonstrate that the coordinated recruitment of specific interneuron subtypes in a biologically-inspired model architecture can transform a global reinforcer into a training signal that guides local plasticity.

## Discussion

We investigated how local interneurons integrate global reinforcement signals into local cortical computation. By leveraging multiple sensory modalities, we were able to discriminate between local and global computational principles of cortical functions. We found that acetylcholine (ACh) inputs to the cortex convey a global reinforcement prediction signal, independent of sensory modality. VIP interneurons (VIP-INs), primary recipients of cholinergic inputs in cortex, encode similar reinforcement prediction signals, but only in the absence of local cortical engagement. The association between local sensory coding and reward prediction signals led to an overall suppression of VIP-INs. This inhibition is likely driven by a subgroup of somatostatin interneurons (SST-INs) which act as a feedback inhibitor, fine-tuning VIP-IN activity to align with local circuit needs. This functional subgroup of SST-INs is characterized by a delayed response to rewards and increased activity for reward-predictive cues. Integrating these mechanisms into a computational model, we propose a circuit-based framework explaining how the ACh-VIP-SST microcircuit could gate learning and enable adaptive learning of local features based on global reinforcement feedback.

### A framework for reinforcement learning in cortex

Understanding the computational principles that govern cortical function remains a formidable challenge in neuroscience^11,70,71^. Traditional models, with their emphasis on feedforward architectures and Hebbian plasticity, fall short of encapsulating the adaptive processes underlying reward-based associative learning in the cortex. The introduction of deep reinforcement learning into the computational neuroscience toolkit offers a fresh perspective on this problem. Yet the application of advanced algorithms (e.g. backpropagation, stochastic gradient descent) to model brain function highlights the gap between computational prowess and the nuanced, biologically-feasible mechanisms of learning and adaptation within the cortex^11,72^. Recent perspectives advocate for a road map wherein system neurosciences would focus on identifying canonical “objective functions, learning rules and architectures” instead of “explicit characterization of neural computation”^11^. Here, while originally looking for behavioral rules for VIP-IN recruitment (**Figure 1 and 2**), we uncovered a canonical architecture and computation able to support reinforcement learning in the cortex. Our observation that neuromodulation inputs on a canonical interneuron motif, known to gate plasticity, offers a compelling and falsifiable framework for reinforcement learning and credit assignment in the cortex.

### Neuromodulation provides a global teaching signal

Neuromodulator inputs to the cortex convey information about both the internal state and environmental feedback. Among the different neuromodulators targeting cortex, basal forebrain acetylcholine strongly drives plasticity of principal cells^63,73,74^. Acetylcholine release in cortex is classically associated with arousal and locomotion^49,50,75,76^, but new lines of evidence link fast cholinergic responses to reinforcement-related signals^57,61,62^. Here we focused on such cholinergic-mediated reinforcement feedback and showed that they provide a global, sensory modality-independent reward prediction coding to the cortex (**Figure 5 and S9**). We identified an interneuron microcircuit that integrates these reinforcement prediction signals and that could guide local plasticity and support associative learning (**Figure 8**). We suggest that our observations could be extended to other modes of recruitment of the cholinergic system, other neuromodulators, or other local cell types expressing neuromodulator receptors. For instance, norepinephrine or serotonergic inputs are likely to provide similar modulation of cortical activity, but the specifics remain to be explored, both in terms of signals being encoded as well as the cortical targets^44,50,77,78^. Conversely, other cell types than VIP-INs are impacted by acetylcholine or other neuromodulators in cortex and are likely to participate in this computation^79,80^.

### Transient disinhibition by VIP interneurons as a canonical learning rule

Interneuron activity is modulated during reinforcement learning, throughout the cortex and across a variety of behavioral tasks. In addition to their role in shaping tuning properties of local principal cells^81^, VIP- and SST-IN activity also reflect the engagement of a canonical inhibitory motif that transforms a global reinforcement signal into local plasticity rules. In this model, VIP-INs are transiently recruited to gate dendritic plasticity mechanisms through disinhibition. Our data support an inhibitory feedback mechanism, mediated by a subtype of SST-INs, which would locally close this plasticity window upon learning (**Figure 7**).

This framework, combining learning-dependent feedforward disinhibition and feedback inhibition has the potential to reconcile, or link together a swath of seemingly contradictory observations. For instance, VIP-INs in the visual cortex can show small reward- and visual-related responses in animals already trained on a visual discrimination task ^29^. This absence of VIP-IN recruitment in the visual cortex parallels our findings in the auditory cortex in expert animals trained on an auditory task (**Figure 1**, see also Ren et al. 2022 for an equivalent observation in motor cortex during motor learning task). In addition, our work supports a learning-dependent recruitment of VIP-INs by reward (see also Krabbe et al., 2018; Ramamurthy et al., 2023). This explains how VIP-INs in naive animals respond more to reward^13,16,17^ than VIP-INs in well trained animals^12,15,29^. Ramamurthy et al., 2023, reported a stronger inhibition of the later reward responses of VIP-INs with learning (see also **Figure 1**) which would align with the delayed recruitment of SSTp-INs as we are now reporting (**Figure S9**).

Several studies and reviews have focused on the heterogeneity of SST-IN activity during learning^29,80,82–88^. Among them, work in the visual cortex shows decreased responses from upper-layer SST-IN in a visual discrimination task^84^. During a motor learning task, a similar bidirectional modulation of distinct SST-INs could be observed in the motor cortex^82^. The feedback inhibition from SST to VIP interneuron we are suggesting here could also parallel a similar feedback inhibition principle found between SST Martinotti cells and principal cells^81^. Overall, these learning- and cell type-dependent VIP-IN and SST-IN dynamics, independently observed across different brain areas and tasks, could be part of a cortex-wide canonical motif gating learning based on top-down feedback.

### A learning architecture supported by genetically-identifiable interneuron subtypes

Until recently, probing the role of interneuron subtype diversity for cortical computation was only accessible by investigating anatomical (morphology, layer location) or electrophysiological correlates. Our study based on genetically identified interneuron groups does not query subgroups of VIP-INs or SST-INs. Nevertheless, we empirically identified two subtypes of SST-INs based on their response to reward which correlated with their layer location (**Figure 6**). New tools arise daily to specifically target subtypes of VIP- or SST-IN using genetic markers. It remains to be seen whether those markers will be useful or could be matched to specific computations as the ones we have identified. In particular, VIP interneuron subgroups have been studied in the hippocampus through the expression of calretinin (CR) or cholecystokinin (CCK). The CR VIP cells are primarily bipolar in shape and primarily target SST cells synaptically, while the CCK cells are classified within the perisomatic basket cell group, capable of innervating both other interneurons and pyramidal cells^46,89,90^. Genetic markers for SST-diversity also exist and have been recently leveraged to study the role of genetically defined SST-IN for learning^87,91^. Recent transcriptomic data from the Allen Brain Institute has identified other subgroups based on different genetically encoded markers. Importantly, VIP and SST interneuron diversity appear highly conserved across brain areas and species suggesting that it could support a specific canonical computation^21,47,92^ such as the one we are proposing here. These new markers would crucially allow us to test the architecture developed in our model (see also methods). In particular, both the connectivity between SST-IN subtypes as well as the inhibitory plasticity^43,80,93^ need to be further explored.

### Predictive processing and credit assignment

Predictive processing is a key feature of the cortical function^71^. Sensory prediction error is observed throughout the cortex^28,94^ and can be modeled using interneuron architecture^95,96^. Our work investigates reinforcement predictive processing and credit assignment in the cortex. While we focused on cholinergic feedback and interneuron motifs to gate plasticity locally, other classes of models have previously addressed this problem using deep reinforcement learning approaches. Integration of reinforcement or top-down feedback and prediction error computation is observed directly at the level of principal cell dendrites^97–99^. Dendritic integration, together with recurrent connection through the cortical hierarchy can be used as substrate for the backpropagation algorithm^69,100,101^. The critical role of SST-IN in controlling dendritic plasticity could bridge these dendritic-dependent models with the interneuron-dependent mechanisms we described here. Our work provides a comprehensive framework for credit assignment in cortex where interneurons play a critical role in integrating global reinforcement feedback into local cortical computation. This framework also provides new perspectives to understand miscalibrated expectations observed in various neuropathologies.

## Materials and Methods

### Animal breeding

Recordings were obtained from adult mice (6–24 weeks of age) of both sexes from the following strains: VIP-Cre, SST-Cre, Chat-Cre and Ai14 (Vip^tm.1^(cre)^Zjh^/J, Sst^tm2.1^(cre)^Zjh^/J, Chat^tm1^(cre)^Lowl^/J, B6.Cg-Gt(ROSA)26Sor^tm9^(CAG–tdTomato)^Hze^/J, The Jackson Laboratory). The water intake was limited to 0.8-1 ml/day following during behavioral training. The mice were provided with unrestricted access to food. Their body weight was monitored biweekly to ensure it remained above 85% of their initial weight. The experimental procedures were conducted in accordance with the guidelines established by the Institutional Animal Care and Use Committee of the Washington University in St. Louis and the Cold Spring Harbor Laboratory and The Animal Care and Experimentation Committee of the Institute of Experimental Medicine of the Hungarian Academy of Sciences. These protocols adhere to the regulations of the EU, Hungarian-, and US National Institutes of Health (NIH), with the reference number: PEI/001/194-4/2014 and PE/EA/54-2/2019.

### Animal surgery

Fiber photometry and 2P-AOD microscopy recordings were performed after similar surgical procedures. Animals were anesthetized with isoflurane (0.5-1.5%) or intraperitoneal injections of a combination of fentanyl, midazolam and medetomidine (0.05/5/0.5 mg/kg, respectively). Mice were positioned in a stereotaxic frame. Local anesthesia (lidocaine or ropivacaine) was administered by subcutaneous injection at the incision site. A small cranial window was then drilled above the targeted brain areas (see below for coordinates). We used a glass micropipette (10-20 μm tip diameter) to deliver the selected virus (see below for virus information. For fiber photometry recordings, a 400 μm fiber (0.48NA, Doric lenses) was then implanted 200 μm above the targeted area. For the 2P-AOD recordings, a dual coverslip was positioned at the location of craniotomy. Both the fiber and the coverslip were then secured in place using a combination of Metabond, light-cured Vitrebond and dental acrylic. All surgery was conducted under aseptic conditions and a heating pad was used to maintain their body temperature. For the following 5 days post surgery, an analgesic was administered daily by intraperitoneal injection (ketoprofen or meloxicam). The mice were then housed for a period of 2 weeks with free access to water and food to allow for recovery and viral expression.

| injection sites | antero-posterior (mm) | medio-lateral (mm) | dorso-ventral (mm) |
| --- | --- | --- | --- |
| auditory cortex | -2.0 | 4.5 | 0.7 |
| basal forebrain | -1.2 | 1.8 | 4 |
| parietal cortex | -2.0 | 1.7 | 0.3 |
| prefrontal cortex | 1.7 | 0.5 | 1 |
| visual cortex | -3.0 | 2.2 | 0.5 |

| virus type | provider | reference |
| --- | --- | --- |
| AAV9.CAG.Flex.GCaMP6f (VIP-cre) | addgene | 100835 |
| AAV9.CAG.Flex.jGCaMP8f (VIP-cre) | addgene | 162382 |
| AAV9.syn.Flex.jGCaMP7f (SST-cre) | addgene | 104488 |
| AAV9.syn.Flex.jGCaMP8f (SST-cre) | addgene | 162379 |
| AAV5.Ef1a.DIO.SwiChRca.TS.EYFP | addgene | 55631 |
| AAV9-hSyn-ACh4.3 | WZ Biosciences | n.a. |
| AAV9-hSyn-ACh4.3mut | WZ Biosciences | n.a. |
| AAV9-hSyn-DIO-ACh4.3 | WZ Biosciences | n.a. |

### Histology

Injection and fiber implantation sites were checked using histological procedures (see Figure S1). Briefly, animals were euthanized using pentobarbital (FATAL-Plus, 150 mg/ml) and an intracardiac injection of paraformaldehyde was quickly performed in order to maintain and fix the brain tissue. After extraction of the brain, 60 μm slices were obtained and directly mounted on a glass slide together with a DAPI staining. We used a Leica THUNDER microscope to acquire fluorescence images at the injection/implantation site.

### Animal behavior

Mice were water-restricted (1 ml of water per 24 hours) between 3 to 7 days prior to the first training session and throughout training. Mice had free access to 3% citric acid water over weekends^102^. Before training the animals on the different Pavlovian conditioning task, mice were habituated to the head fixation apparatus and received and collected random 5 µl water reward for 5 days. During this period, the auditory response in the auditory cortex was evaluated using an auditory tuning protocol consisting of the random delivery of 200 ms pure tones ranging from 4 kHz to 20 kHz, upward and downward chirps (from 4 kHz to 20 kHz) and white noise, repeated 5 times. The same auditory tuning protocol was presented on the last day of training of the auditory task (see Figure 2c). Recordings from the auditory cortex that did not show any sound responses were discarded and the animals excluded from this study. After this habituation phase, animals were trained and recorded daily on a Pavlovian, trace conditioning task. The task required animals to associate a sensory stimulus with the delayed delivery of a water reward. This conditional stimulus was presented on 40% of the trials and was followed 80% of the time by a 5 µl reward droplet. The delay between the conditional stimulus and the reward delivery was set to 1 s for the visual and auditory task and to 2 s for the olfactory task. 2P-AOD microscopy recordings also used a 2 s delay. For control trials, a neutral stimulus, an uncued reward, or a blank trial with no cue or reward delivered were presented 30%, 15%, and 15% of the time, respectively. The auditory cues consisted of upward (4kHz to 20kHz) or downward (20kHz to 4kH) chirps generated using ‘chirp’ Matlab function. The visual cues consisted of the activation of an LED positioned on the left or on the right side of the animals. The olfactory task in Figure S8 used a custom made delivery system to present the odors at a constant flow rate of 1 l/min, with in-line filters (Whatman 6823-1327) containing 20 µl of 1% isoamyl acetate (SigmaAldrich W205508) or Ethyl tiglate (SigmaAldrich W246000). The conditional cue identity was selected at random on the first day of training and results were pulled together independently of the cue identity. Animals were trained on one or a succession of these tasks using different sensory modalities.

The go-nogo task used in Figure S5-6 were performed as in Szadai et al. 2022. All behavioral protocols were controlled using Bpod (SanWorks) and Matlab routines (https://github.com/QuentinNeuro).

### Fiber photometry

Fiber photometry data were collected and analyzed using a custom-made photometry setup and Matlab routines (see also Fluorescence analysis section below). We used 470 nm (M470F3, Thorlabs) and 405 nm (M405FP1, Thorlabs) LED sources coupled to an optic fiber (M75L01, Thorlabs) and collimation lens (F240FC-A) for fluorescence excitation. The excitation lights were delivered to the cannula implanted on the head of the animal using a second collimation lens (F240FC-A, Thorlabs) coupled to a 400 μm, high NA, low autofluorescence optic fiber (FP400URT, custom made, Thorlabs). The emission light was collected using the same optic fiber and directed to a Newport 2151 photoreceiver using a focusing lens (ACL2541U-A, Thorlabs). Excitation filters (ET470/24 M, ET405/10M), emission filters (ET519/26M), and dichroic mirrors (T495LPXR, T425LPXR) were purchased from Chroma Technology. The 470-nm and 405-nm excitation lights were respectively amplitude-modulated at a frequency of 211 Hz and 531 Hz, with a max power of 40 μW, using an LED driver (LEDD1B, Thorlabs) controlled through a National Instrument DAQ (NI USB-6341). The modulated data acquired from the photoreceiver were decoded as in Lerner et al., 2015.

### 2-photon acousto-optical deflector (2P-AOD) imaging

We used the Femto3D Atlas multiphoton microscope (Femtonics) to record single VIP and SST interneuron activity and acetylcholine release surrounding VIP interneurons. This system uses acousto-optical deflectors (AOD) for ultrafast two-photon fluorescence excitation (920 nm, MaiTai HP, SpectraPhysics or a Coherent Chameleon Ultra II, Coherent). AODs use chirped sinusoidal acoustic waves in piezoelectric crystals for z and xy plane scanning. A second set of deflectors and two automatic laser beam stabilizers (BeamStab, Femtonics) were used for drift correction. We used a 16x N16XLWD-PF Nikon objective for excitation light delivery to, and emission light collection from the window implanted above the target area. A dichroic mirror (700dcrxu, Chroma Technology) was used to separate emission from excitation wavelengths, and fluorescent signals were acquired using GaAsP photomultipliers (H10770PA-40, Hamamatsu). For the acetylcholine release and tdTomato-expressing VIP interneuron recordings, GRAB-ACh and tdTomato signals were separated using a dichroic mirror (t600lpxr, Chroma Technology) and emission filters (ET520/60m, ET650/100m, Chroma Technology).

The Femto3D Atlas multiphoton microscope allows for ultrafast scanning of regions of interest (ROI) selected in a 3D volume. Before training, we acquired a 500*\**500*\**500 μm (x*y*z) (0.5-1 pixels/μm x, y resolution and 10 μm z-steps) z-stack to map the interneuron locations in the target brain area. Individual cells were marked using the acquisition software (MES, Femtonics) and a squared ROI encompassing the center of the cells were then automatically generated (FoldedFrame imaging method). These ROI were then scanned at 10-30 Hz, using a 1 - 2 pixels/μm resolution.

### Optogenetic manipulation

For Figure S6, we implanted a 200 um optic fiber implant (FT200EMT, Thorlabs) 0.5 mm above the injection site and with a 45 degree angle to allow for imaging in the dorsal cortex. The stimulation consisted of a 3 seconds continuous pulse, starting at cue delivery using a 470 nm laser (PSU-III-LED, Changchun New Industries Optoelectronics Technology Ltd., 10-18 mW/mm2). The opsin was allowed to turn back off using a long inter-trial interval that exceeded 40 seconds.

### Behavioral analysis

We compared data from the first day of training (early in the text) to the last days (late in the text, between 1 and 3 sessions per animal). The licking rate used to measure the behavior performance was computed on all trials. Only trials where the animals collected the reward droplets were included for cued reward (CR) and uncued reward (UR) trials. For conditional stimulus (CS) and neutral stimulus (NS) neuronal data analysis late in training, trials were filtered based on the anticipatory licking (CS>1Hz, NS<1Hz) behavior to decrease within-session variability.

In Figure S5, we reanalyzed recordings from^16^. We further divided the recordings based on mouse behavior. Animals were considered as increasing their performance during the session if they showed an increase in hit rate of more than 5 percentage points between the early (first 20 trials) and late part of the recording sessions (next 60 trials).

### Fluorescence analysis

Data from fiber photometry and 2P-AOD imaging were analyzed using the same Matlab routines (https://github.com/QuentinNeuro). Single trial fluorescent traces were z-scored or DFF using the mean and standard deviation values measured during a one second window preceding the cue delivery. To mitigate single trial variation in baseline, we used a 5 trial moving median average of the mean and standard deviation baseline values before normalizing the data. The average, or maximum amplitudes were then computed on the average or single trial traces for the cue (that included cue and delay) and reward period (2 sec window from reward delivery). The fluorescent value directly preceding the cue or reward period was then subtracted to obtain a relative change in fluorescence.

Single VIP-IN data from 2P-AOD recordings were separated as responding or not responding to reward after normalizing the DFF reward responses by the standard deviation of the baseline. VIP-INs were considered reward responsive if their reward responses exceeded 3 baseline standard deviations.

Isosbestic controls were realized on a subset of animals throughout the study, to control the contribution of hemodynamics to the changes in fluorescence measured with fiber photometry. 470-nm and 405-nm fluorescent data were normalized using the equation in Figure S1 which was compared to DFF 470-nm fluorescent data. For the acetylcholine sensor data, we used a mutated version of the receptor as control since the isosbestic point is undefined.

In Figure 5, we computed a residual fluorescence metric for the cholinergic recordings to account for the slower dynamics of the signals. The residual fluorescence for the cue period was calculated by subtracting the average NS fluorescence to the CS fluorescence, thus revealing the difference between CS and NS cholinergic activity. The residual fluorescence for the reward period was calculated by subtracting the non-rewarded CS trials to the reward cued trials, thus removing the cue dynamics from the reward-related changes in cholinergic activity.

### Statistical analysis

Data are represented as mean ± SEM. Boxplots in Figure 1 and 6 were generated using the boxplot Matlab function and show the median (centerline), 25 and 75th percentile (box), minimum and maximum values (whisker) as well as outlier data (crosses). Student’s t-tests were computed using sigmaStat and Matlab functions after checking for the normality and the homoscedasticity of the distributions to compare. Otherwise, we used non-parametric Wilcoxon signed-rank (same population, WSRT in the text) or Mann-Whitney U (different population, MWT) tests. Single trial (Figure 8) or single cell (Figure 3) distributions were compared using a Kolmogorov–Smirnov test (KS). Cumulative distributions shown in Figure 8 were obtained after randomly selecting 100 trials per animal. All cells from all 3 recordings were included in the cumulative distribution in Figure 3. All data were considered significantly different when p<0.05.

### Computational model

We used a rate-based mean-field model with an excitatory pyramidal cell (PC), an inhibitory vasoactive intestinal peptide-expressing interneuron (VIP-IN), as well as two types of somatostatin-expressing interneurons (SSTn-IN and SSTp-IN).

The PC is divided into two coupled compartments, representing the soma and the dendrites, respectively. The dynamics of the firing rate of the somatic compartment, *r*_PC_, obeys ^104^:

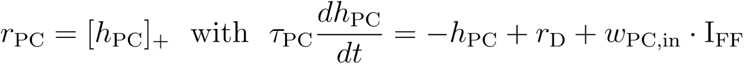

Firing rates are rectified to ensure positivity, […]_+_.*τ*_PC_ is the time constant of the excitatory neuron (*τ*_PC_ = 20 ms), the weight *w*_PC,in_ denotes the connection strength between the sensory feedforward input and the PC.

Similarly, the activity at the dendrite, *r*_D_, evolves according to

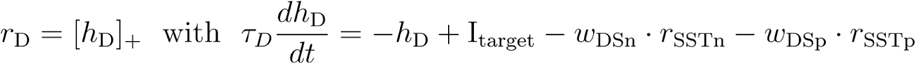

*τ*_D_ is the time constant of the dendritic compartment (*τ*_D_ = 50 ms), the weights *w*_DSn_ and *w*_DSp_ denote the dendritic inhibition provided by the SSTn-IN and the SSTp-IN, respectively (see below). I_target_ represents a sensory target, potentially arriving from higher-order areas. The firing rate dynamics of each interneuron (SSTp, SSTn, VIP) is modeled by a rectified, linear differential equation:

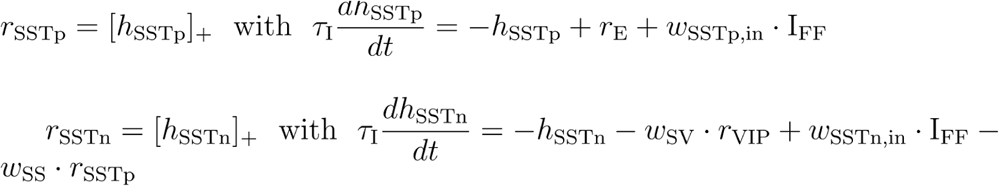

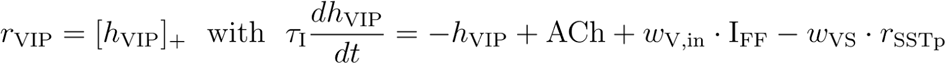

*τ*_I_ is the time constant of the inhibitory neurons (*τ*_I_ = 10 ms). The weights *w*_SSTp,in_, *w*_SSTn,in_ and *w*_V,in_ denote the weights between the sensory feedforward input and the respective interneuron (*w*_SSTp,in_ = 0, *w*_SSTn,in_ = 2.5, *w*_V,in_ = 2). The weights *w*_SV_ and *w*_SS_ represent the inhibition strength from the VIP-IN and the SSTp-IN onto the SSTn-IN, respectively, and the weight *w*_VS_ denotes the inhibition from the SSTp-IN to the VIP-IN (*w*_SV_ = 0.5, *w*_SS_ = 0.9, *w*_VS_ see below). While SST-to-SST connections have been reported to be largely missing, a study by Jiang et al., 2015 found those connections between subgroups of SST-INs. In our model, this synaptic connection ensures that SSTn-IN activity decreases from the early to late phases of learning. However, other mechanisms are conceivable. For instance, we do not model PV interneurons in our network. PV interneurons most likely will also receive direct local excitation from the PC and may increase its activity upon learning the cue-outcome association. Hence, another source of inhibition, like from PV interneurons, may also explain the decrease in SSTn-IN activity without the need for a direct connection from SSTp- to SSTn-INs.

In addition to the sensory feedforward input I_FF_, the VIP-IN also receives acetylcholine (ACh). According to the experimental data, we assume that ACh is initially low before reaching a steady state at the cue location. For the sake of simplicity, we model ACh as a sigmoid function:

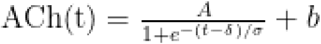

*A* is the amplitude (strength of ACh), *t* denotes the time, δ and σ characterize the shift and steepness of the sigmoid function, respectively, and b denotes the baseline ACh level. In the simulations, we use *A* = 2.5, δ = 700, and σ = 20, b = 0.5. Furthermore, the sensory feedforward input I_FF_ is set to 1 in the auditory task and 0.02 in the visual task, while I_target_ is fixed at 2.

*w*_DSp_ was chosen such that when the PC reaches the sensory target, the feedback inhibition from SSTp-IN cancels the sensory target at the dendritic compartment (here *w*_DSp_ = 1). In addition, *w*_DSn_ is set such that the overall dendritic inhibition balances the sensory target at the beginning of learning:

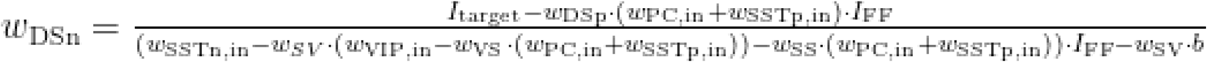

In our model, the learning window closes because the SSTn-IN is eventually released from VIP-IN inhibition. The VIP-IN suppression is shaped by two local processes. First, the VIP-IN receives more inhibition from the SSTp-IN as a consequence of the PC learning the sensory target. This is achieved through changes in *w*_PC,in_ that evolves according to

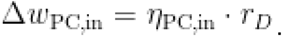

While it is conceivable that the suppression of VIP-IN activity is solely achieved through the increase of SSTp-IN activity, we found that the parameter ranges for which the early/late VIP-IN activity is qualitatively similar to the data are broader with an accompanying mechanism that increases the inhibition onto the VIP-IN. Hence, we assume that the synapse between the SSTp-IN and VIP-IN strengthens through inhibitory Hebbian plasticity^105^:

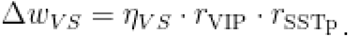

While we chose long-term changes in the SSTp-to-VIP synapse, more transient changes, for instance, neuronal adaptation or short-term facilitation^22,46,106^. To ensure that the PC learns to predict the target before the plasticity window closes, that is, before the SSTn-IN is released from VIP-IN inhibition, the cue-learning must be faster than the changes in the synapses from the SSTp- to the VIP-IN. The learning rates *η*_PC,in_ and η*_VS_* are set to 0.04 and 0.0003, respectively. The initial weights were 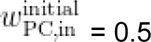 and 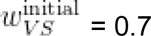. All other weights were fixed.

## Author contributions

Q.C. and A.K. conceived, designed, and interpreted the study. Q.C. performed and analyzed the experiments and wrote the manuscript. Z.S. and B.R. designed, performed and analyzed 2P-AOD experiments. E.T.G. and M.M. performed fiber photometry experiments. L.H., Q.C and R.P.C. designed the model, L.H. constructed and analyzed. A.K. supervised the project and edited the manuscript. All authors provided comments on the manuscript.

## Acknowledgements

We are grateful to Dr. A. Bacci, Dr. J. Cardin and the Kepecs lab members Dr. S. Li and E. Bano for comments on a previous version of this manuscript, as well as Dr. H.J. Pi, M. Cortez and J. Dahan for technical support. This work was supported by National Institute of Mental Health (NIH) grants and BJC investigator fund to A.K., by Medical Research Council (MR/X006107/1), BBSRC (BB/X013340/1) and ERC-UKRI Frontier Research Guarantee Grant (EP/Y027841/1) to R.P.C. and by KFI-2018-00097, VKE-2018-00032, NKP-2017-00001 to B.R.

## Supplementary figures

**Figure S1.**
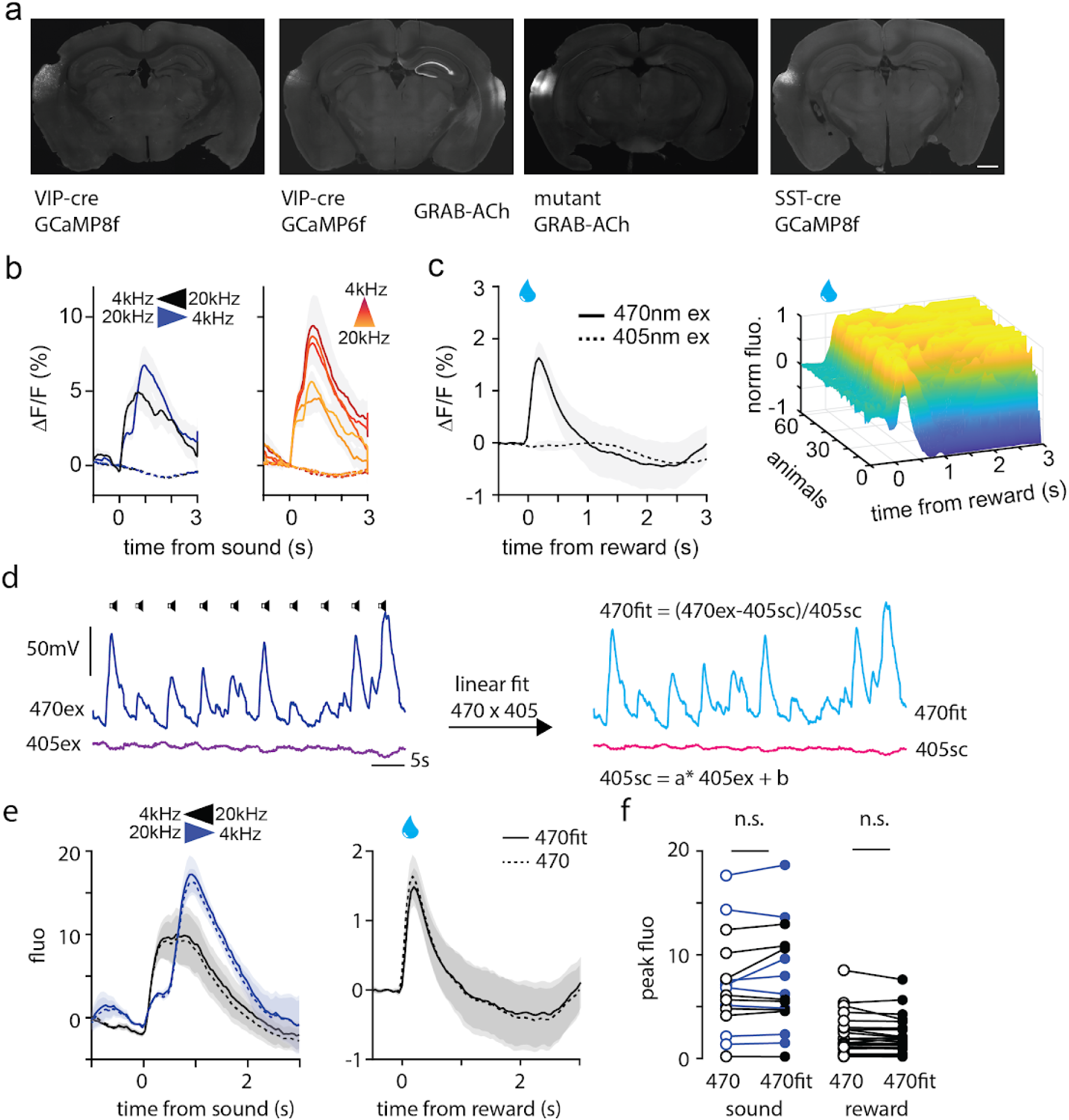
Histology and isosbestic controls for photometry recordings. (a) Example histology images for auditory cortex viral injection and fiber implantation. (b) Chirp (left) and pure tone (right) evoked VIP-IN responses using 470 nm (solid) or 405 nm excitation light (dash). (c) Reward-evoked VIP-IN responses using 470 nm (solid) or 405 nm excitation light (dash) (left). 3D, pseudo-colored plot of all reward-evoked VIP-IN responses recorded in this study (n=63, right). (d) Example fluorescent recording traces from 470 nm and 405 nm excitation lights, before and after fitting procedure (linear fit parameters: 405sc=a*405ex+b, a=0.69, b=-0.05). (e) Chirps and reward evoked VIP-IN responses before and after 405 fit correction. (f) Quantification of the contribution of non-GCaMP dependent fluorescent signals to the sound- (n=8, 5+/-0.3%, p = 0.930 ranked ANOVA) and reward- (n=26, 4+/-3%, p=0.06 WRST) evoked responses of VIP-INs. n.s.: non-significant, scale bar 1 mm.

**Figure S2.**
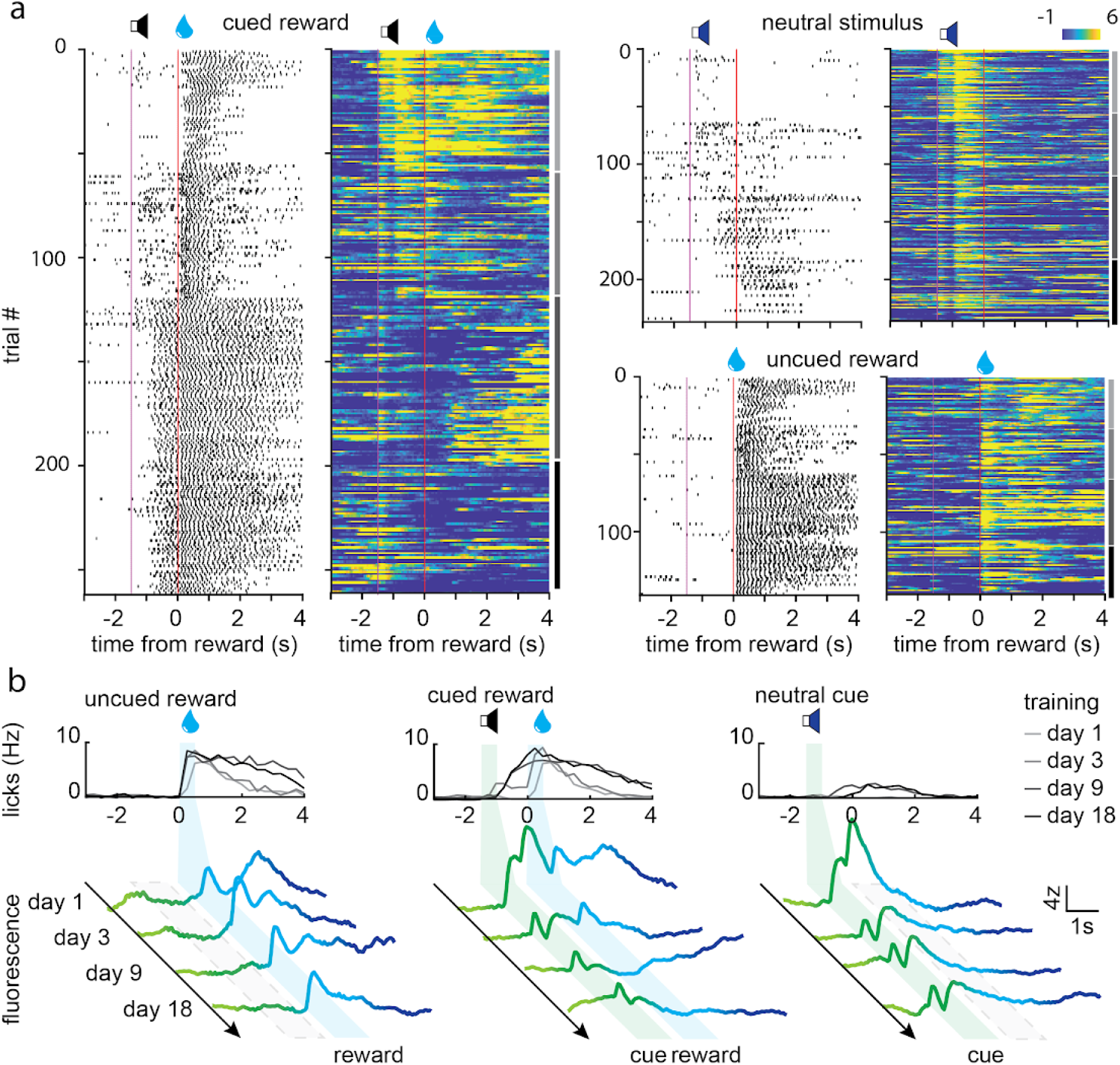
Single trial dynamics of cue and reward responses during auditory task learning. (a) Raster plots showing licking behavior and VIP-IN activity during 4 different days of task learning. (b) Average lick rate and VIP-IN activity during the same training days as in (a).

**Figure S3.**
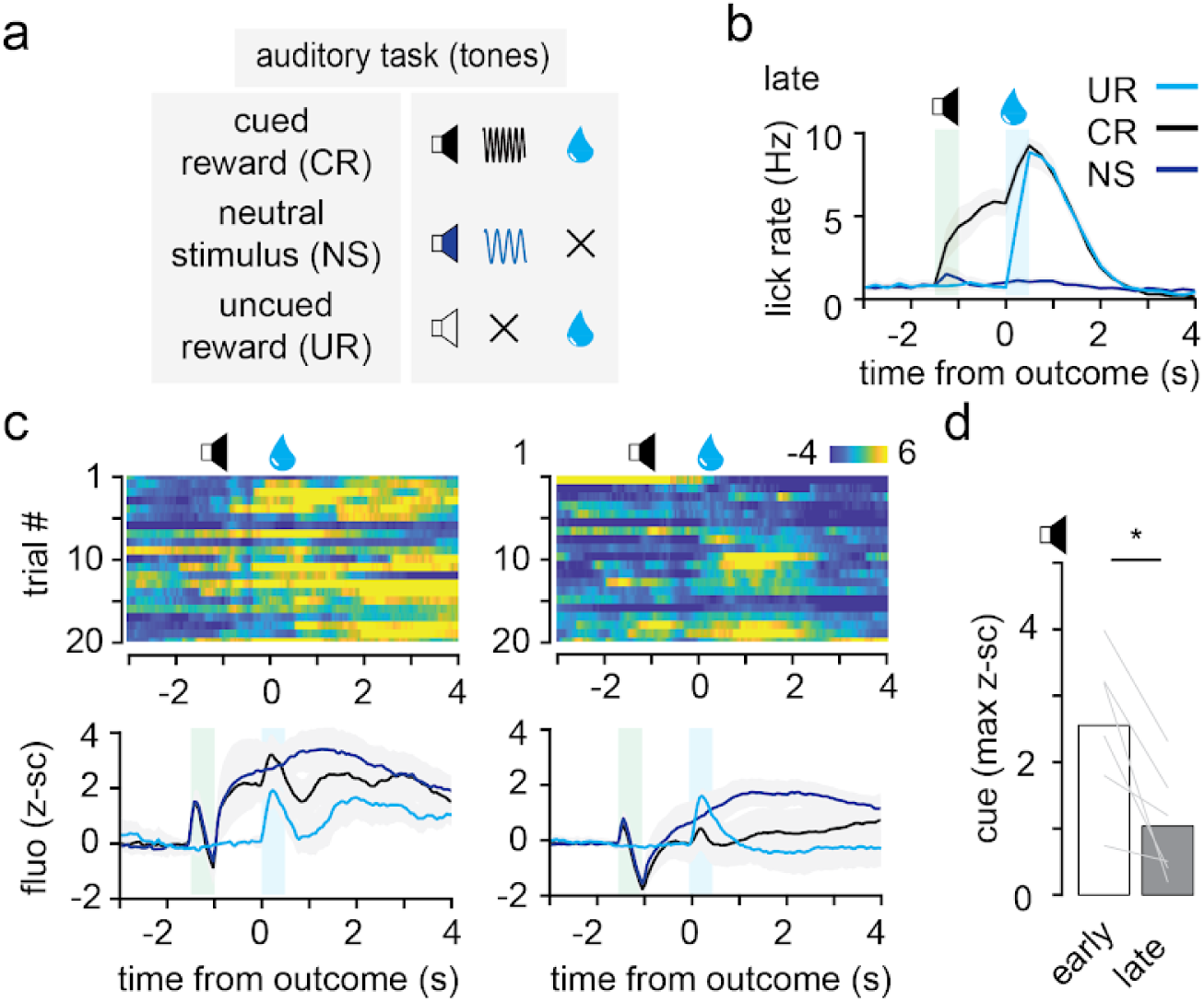
VIP-INs activity during auditory task using pure tones. (a) Auditory task training schedule. (b) Average lick rate late in training (n=6). (c) Pseudo-colored raster plot showing z-scored VIP-IN single trial activity for early (left) and late (right) in training (top). Average VIP-IN activity early (left) and late (right) in training for trial types, colored as in B (n=6). (d) Quantification of the average VIP-IN cue (top) and reward (bottom) responses.

**Figure S4.**
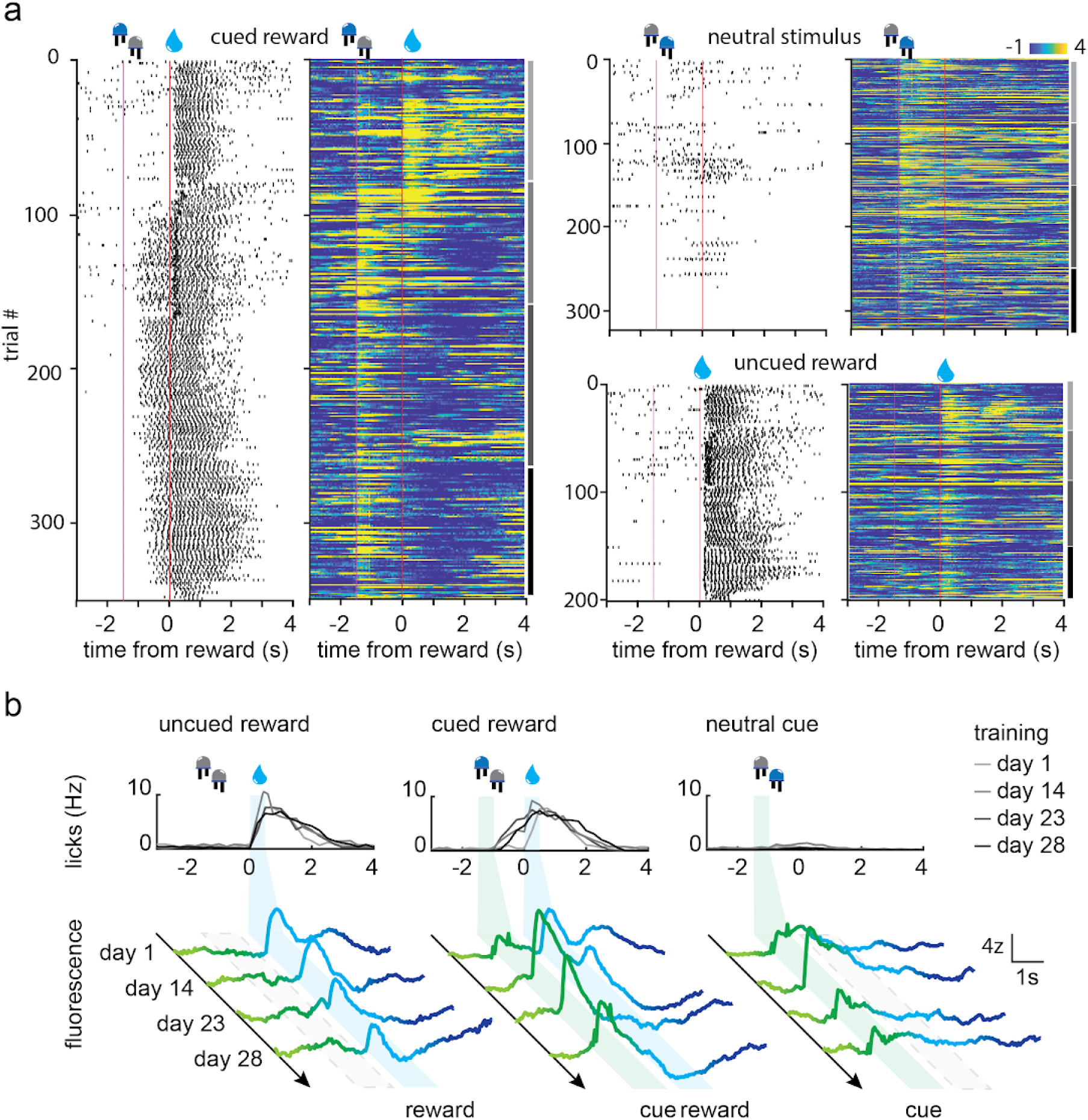
Single trial dynamics of cue and reward responses during visual task learning. (a) Raster plots showing licking behavior and VIP-IN activity during 4 different days of task learning. (b) Average lick rate and VIP-IN activity during the same training days as in (a).

**Figure S5.**
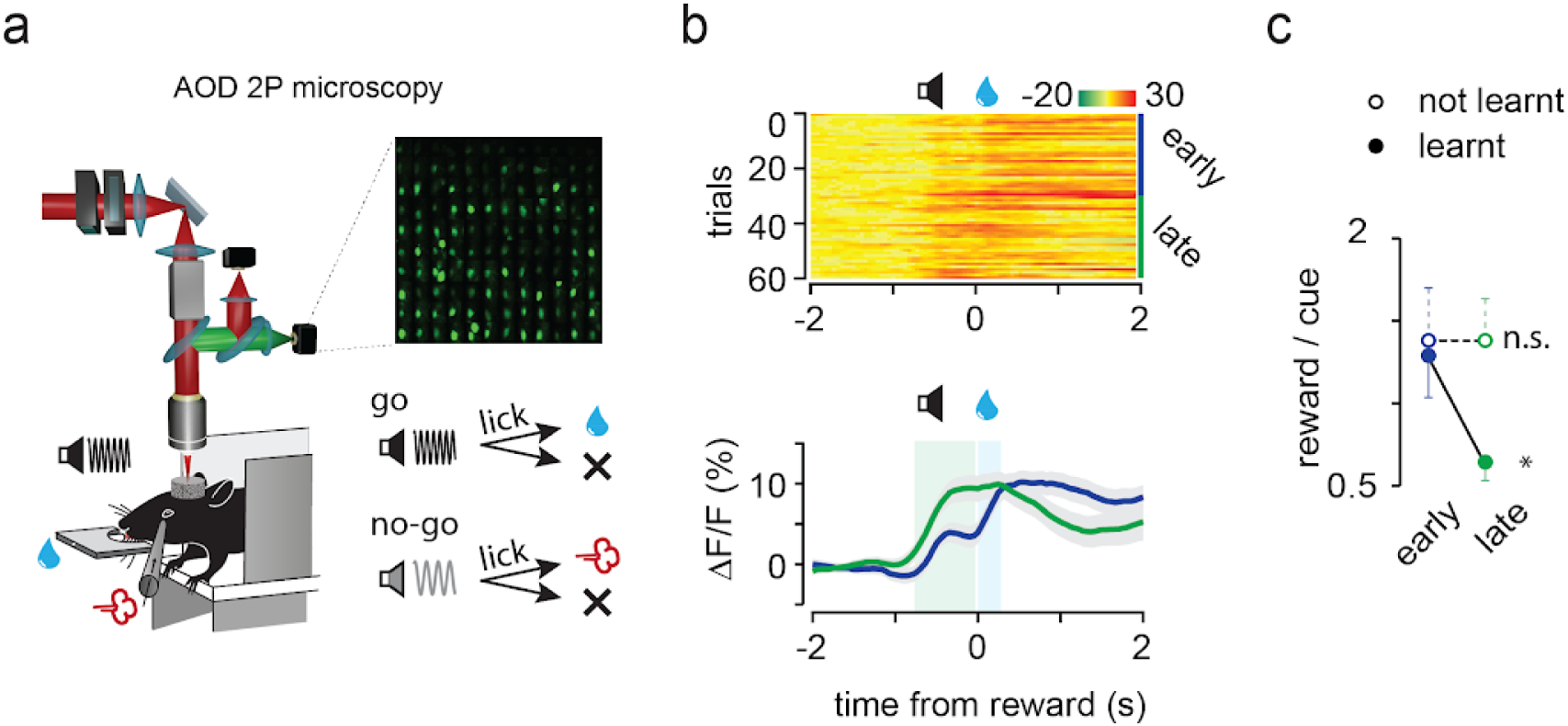
VIP-INs in mPTA show similar learning-related cue coding during a go-nogo task. (a) Diagram of the AOD 2-photons setup and behavior schedule as used in Szadai et al. (b) Top: Raster plot of single VIP-IN activity during learning of the auditory go-nogo task. Bottom: average VIP-IN activity during go trials for animals that showed within-task increase in performance. (c) Quantification of the Reward/Cue VIP-IN response for animals that showed an increase in performance (learnt) vs. not (not learnt) within a single session (n=17).

**Figure S6.**
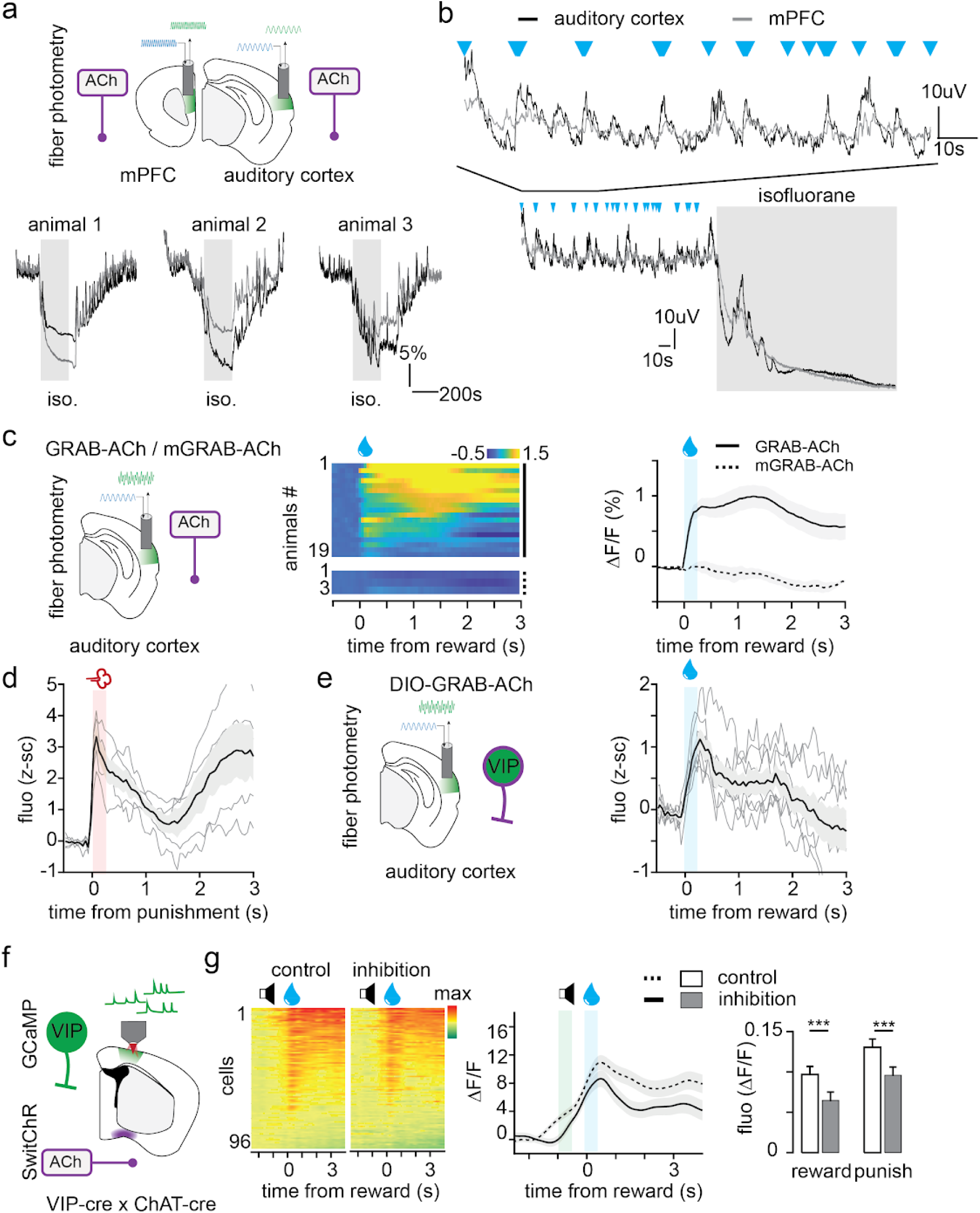
Characterization of cholinergic inputs to cortex. (a-b) GRAB-ACh co-recording in ACx and mPFC showing acetylcholine release dynamics during awake, reward delivery and anesthetized state. (c) Diagram of fiber photometry recordings of ACh release in the auditory cortex (left). Raster (middle) and average (right) uncued reward-related changes in fluorescence from GRAB-ACh sensor (n=19) or from the mutated ACh insensitive version of the sensor (mGRAB-ACh, n=3). (d) Average punishment-mediated GRAB-ACh fluorescence recorded in the auditory cortex using fiber photometry (n=5). (e) Average reward-mediated GRAB-ACh-expressing VIP-IN fluorescence recorded in the auditory cortex using fiber photometry (n=7). (f) Diagram of VIP-IN recording in mPTA while inhibiting cholinergic inputs from the basal forebrain using SwitchR. (g) Raster plot, average traces and quantification of the inhibition in VIP-IN responses to reward and punishment upon inhibition of cholinergic inputs.

**Figure S7.**
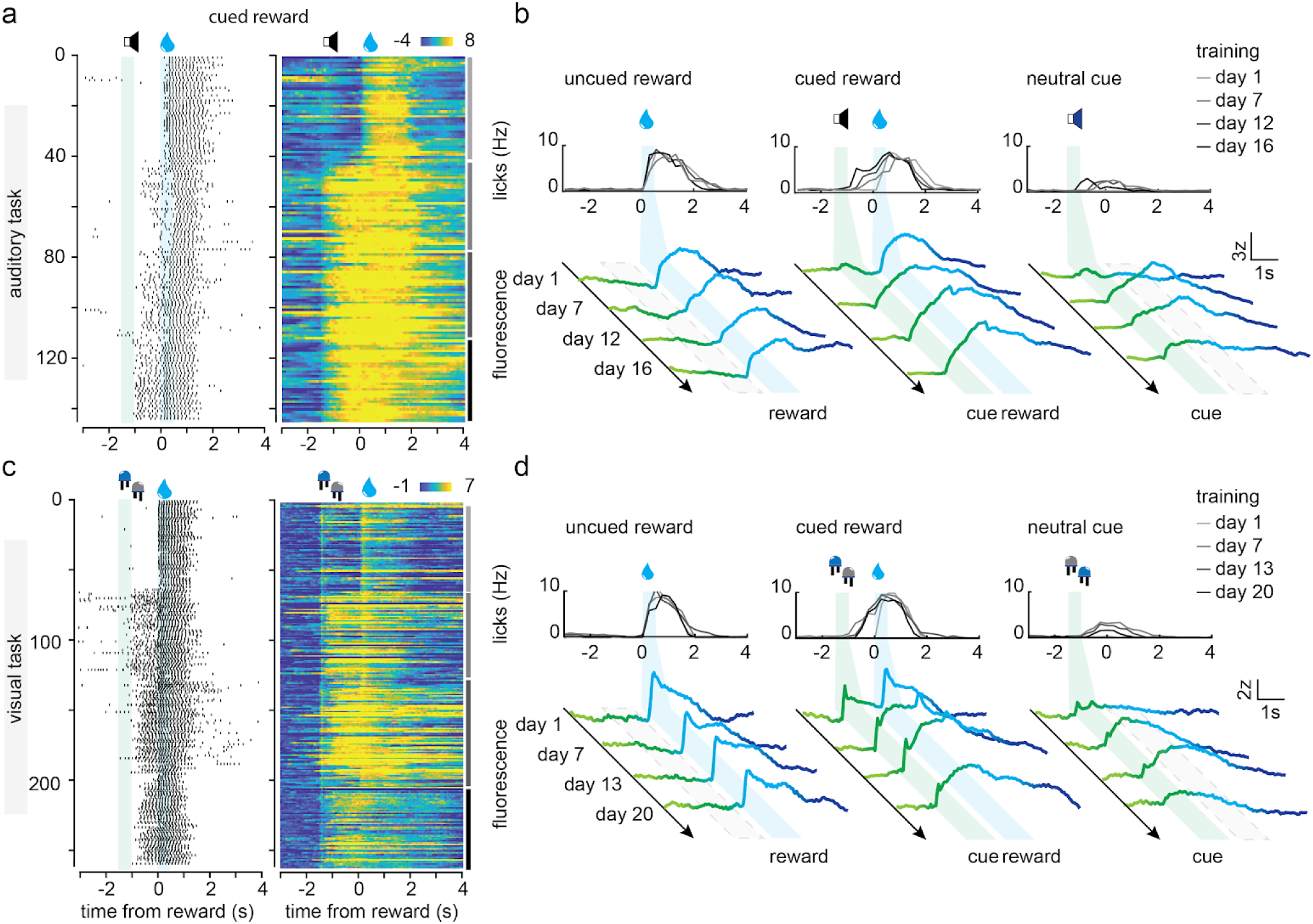
Single trial dynamics of ACh release during the auditory and visual tasks. *(a)* Raster plots showing licking behavior and GRAB-ACh activity during 4 different days of auditory task learning. (b) Average lick rate and GRAB-ACh activity during the same training days as in (a). (c-d) same as (a-b) but for the visual task.

**Figure S8.**
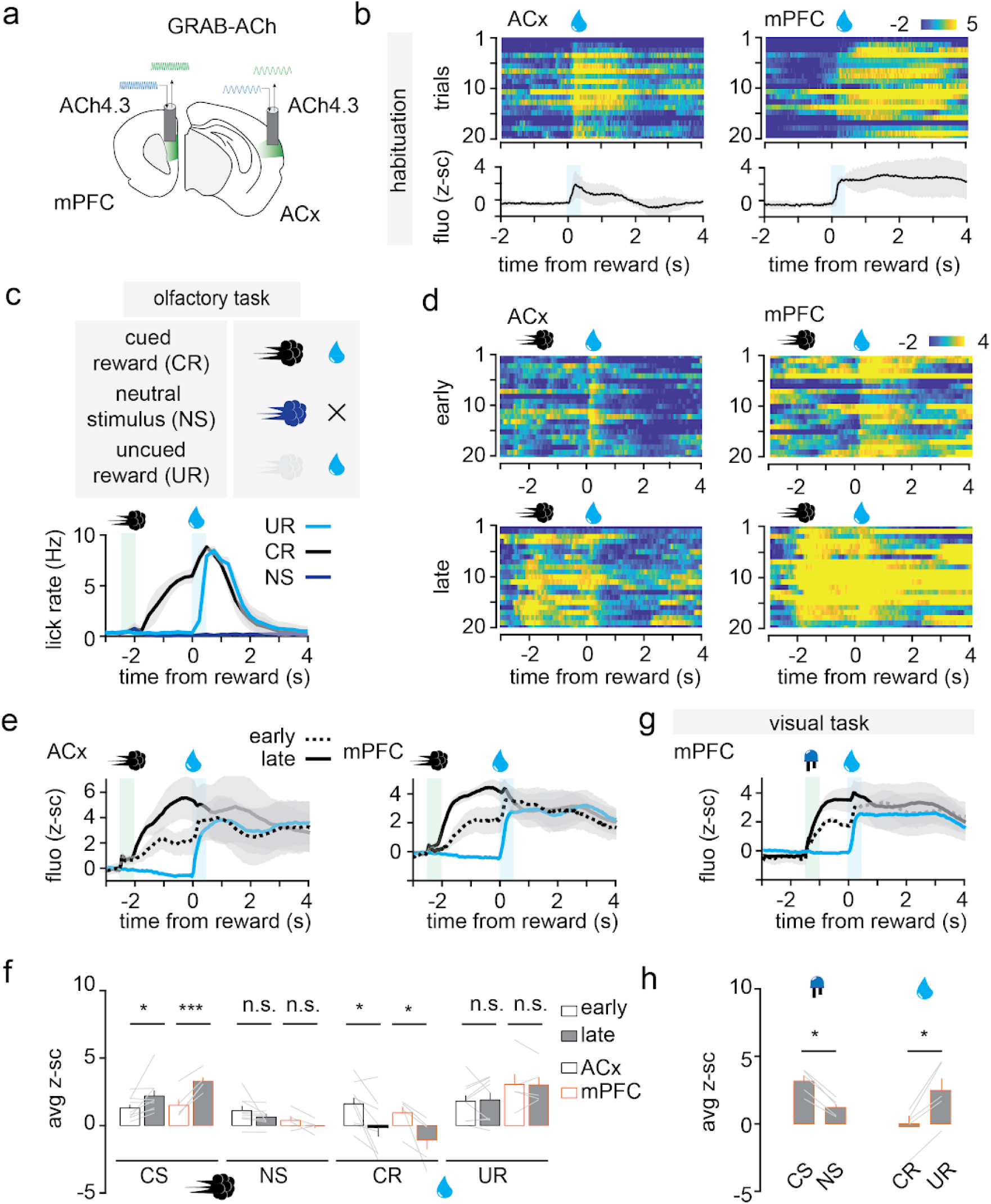
ACh release in ACx and mPFC during conditioning training. (a) Diagram of the GRAB-ACh fiber photometry recordings in mPFC and auditory cortex. (b) Pseudo-colored z-scored fluorescence raster plot for uncued reward trials. (c) Diagram of the olfactory task protocol, and average lick rate after training (n=9). (d) Pseudo-colored z-scored fluorescence raster plot for cued reward trials early and late in training in ACx (n=9) and mPFC (n=5). (e) Average z-scored fluorescence changes in the ACx and mPFC during training of the olfactory task. (f) Quantification of the cue- and reward-related fluorescence in the ACx and mPFC early and late in training of the olfactory task. (g) Average z-scored fluorescence changes in mPFC during the visual task. (h) Quantification of the cue- and reward-related fluorescence in the mPFC late in training of the visual task. CS: Conditional Stimulus, NS: Neutral Stimulus, UR: Uncued Reward, CR: Cued Reward, NS: Neutral Stimulus. n: mice, n.s.: non-significant, *: p<0.05, **: p<0.01, ***: p<0.005.

**Figure S9.**
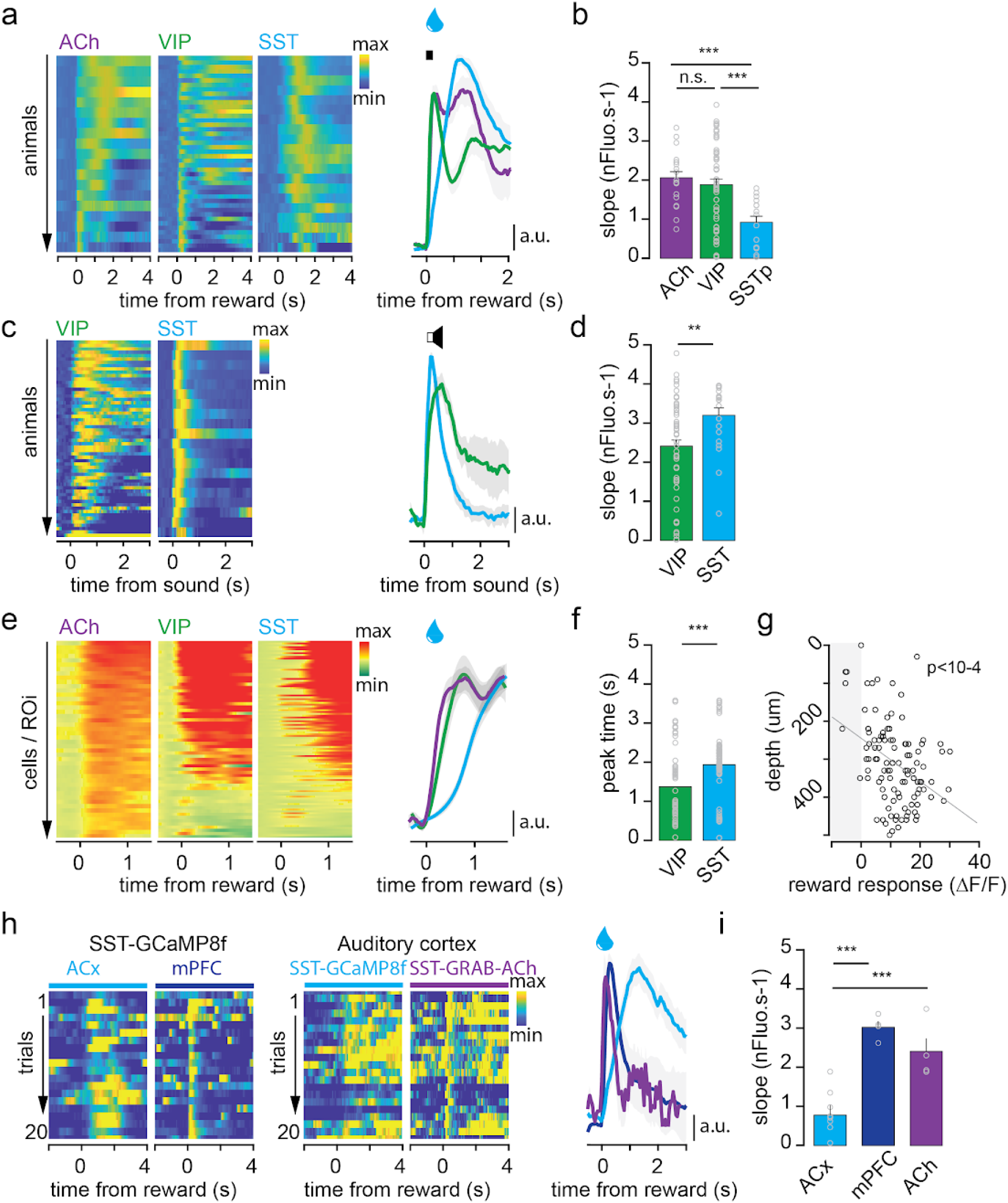
Reward-dependent recruitment of SST-INs is slower than VIP-INs and acetylcholine. *(a)* Photometry recordings of reward-related fluorescence from GRAB-ACh (n=18), GCaMP-expressing VIP-INs (n=55) and SST-INs (n=15) in auditory cortex. (b) Quantification of the reward-evoked slope transients using the time window depicted in (a) (0-0.2s). (c-d) Same as (a-b) but for sound-evoked transients from VIP-IN (n=58) and SST-IN (n=20) recorded in the auditory cortex. (e-f) Same as (a-b) but for single VIP-IN (n=61 cells), SST-IN (n=107 cells) and GRAB-ACh region of interest (ROI) recorded using 2P-AOD microscopy in parietal and visual cortices. (g) Correlation between single SST-IN reward-mediated response and depth (n=107 cells). (h-i) Same as (a-b) but for reward-evoked transients from SST-IN co-recorded in auditory (ACx, n=8) or medial prefrontal cortex (mPFC, n=4, left) or GRAB-ACh-expressing SST-IN in auditory cortex (n=4, right). n: mice, **: p<0.01, ***: p<0.005, n.s.: non-significant.

**Figure S10.**
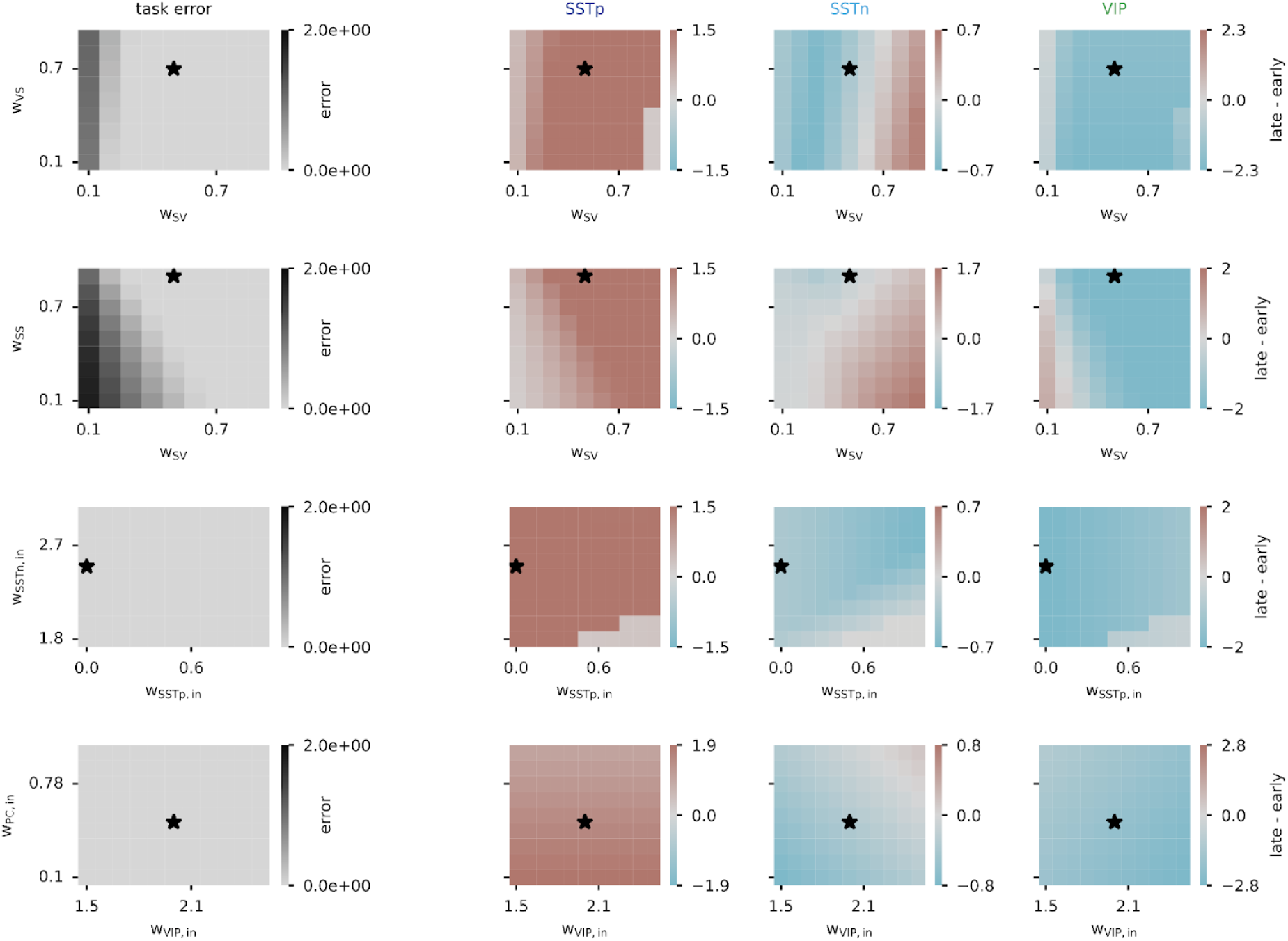
Modeled cue responses from INs early and late in training for the auditory task for different connectivities. Columns, left to right: Task error at the end of training, SSTp-IN, SSTn-IN, VIP-IN. Rows, top to bottom: Difference in cue responses between late and early learning (late - early). Strength of connection between VIP-IN and SSTn-IN (wSV) and between SSTp-IN and VIP-IN (wVS) varied, strength of connection between VIP-IN and SSTn-IN (wSV) and between SSTp-IN and SSTn-IN (wSS) varied, strength of connection between local input and SSTp-IN (wSSTp,in) and between local input and SSTn-IN (wSSTn,in) varied, strength of connection between local input and VIP-IN (wVIP,in) and between local input and the excitatory cell (wPC,in) varied. Star denotes the parameters used in Figure 8.

**Figure S11.**
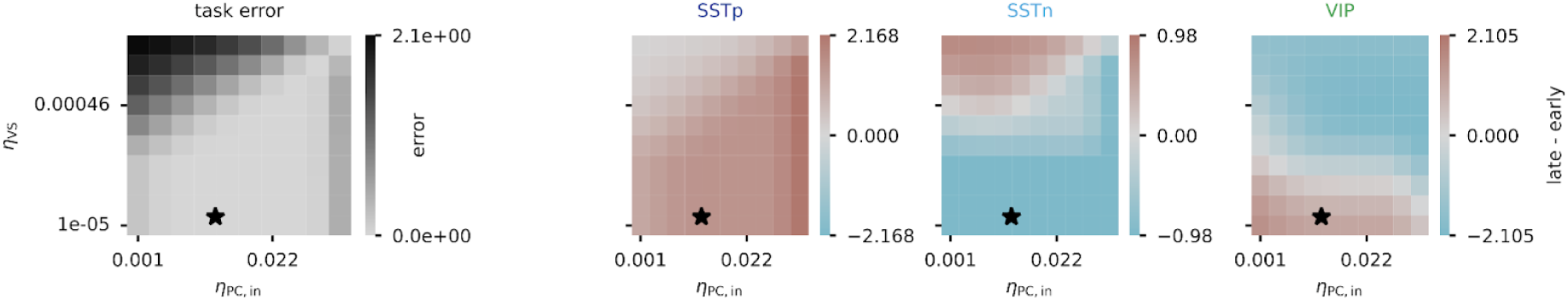
Modeled cue responses from INs early and late in training for the auditory task for different learning rates. Columns, left to right: Task error at the end of training, SSTp-IN, SSTn-IN, VIP-IN. Difference in cue responses between late and early learning (late - early). Learning rate for the connection from the local input to the excitatory cell, and learning rate for the connection from SSTp-IN to VIP-IN are varied. Star denotes the parameters used in Figure 8.

**Figure S12.**
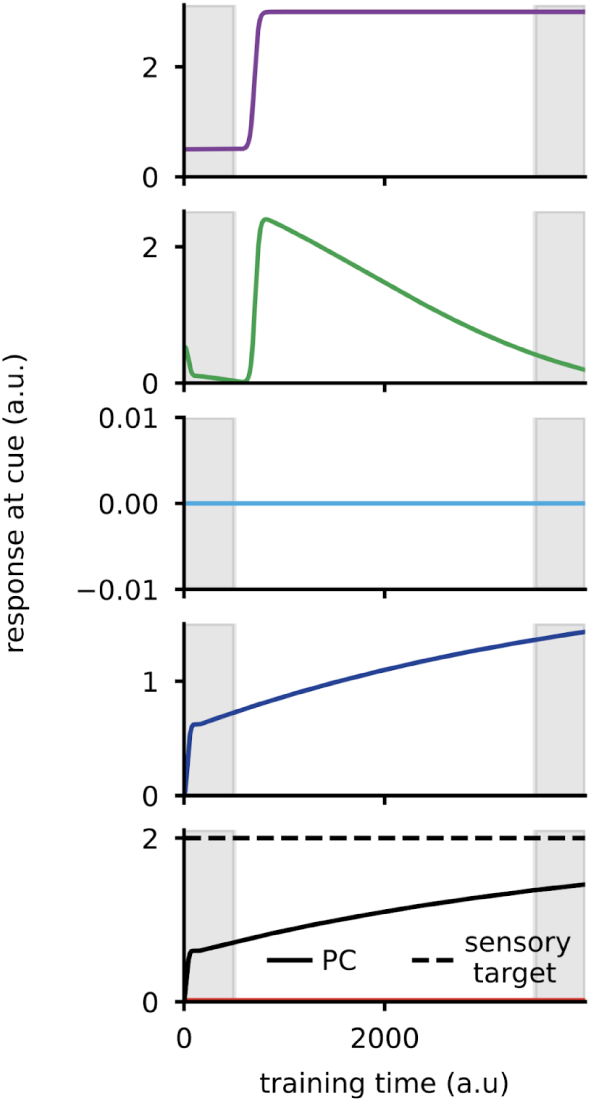
Dynamics of modeled ‘cue’ responses from each neuronal cell type for the visual task. Colors as in Figure 8a. Top to bottom: ACh signal modeled as sigmoid function, VIP-IN, SSTn-IN, SSTp-IN, excitatory cell (black) with sensory target (black dashed) and local sensory input (red). Local sensory input is almost zero.

## Notes

### Competing Interest Statement

Balazs Rozsa is a founder of Femtonics Ltd. and a member of its scientific advisory board

## References

1. Botvinick, M., Wang, J. X., Dabney, W., Miller, K. J. & Kurth-Nelson, Z. Deep Reinforcement Learning and Its Neuroscientific Implications. Neuron 107, 603–616 (2020).

2. Sutton, R. S. & Barto, A. G. Reinforcement Learning, Second Edition: An Introduction. (MIT Press, 2018).

3. Tomov, M. S., Tsividis, P. A., Pouncy, T., Tenenbaum, J. B. & Gershman, S. J. The neural architecture of theory-based reinforcement learning. Neuron 111, 1331–1344.e8 (2023).

4. Bayer, H. M. & Glimcher, P. W. Midbrain Dopamine Neurons Encode a Quantitative Reward Prediction Error Signal. Neuron 47, 129–141 (2005).

5. Doya, K. Reinforcement learning: Computational theory and biological mechanisms. HFSP J. 1, 30–40 (2007).

6. Schultz, W. Dopamine reward prediction-error signalling: a two-component response. Nat. Rev. Neurosci. 17, 183–195 (2016).

7. Schultz, W. Predictive Reward Signal of Dopamine Neurons. J. Neurophysiol. 80, 1–27 (1998).

8. Dayan, P. Dopamine, Reinforcement Learning, and Addiction. Pharmacopsychiatry 42, S56–S65 (2009).

9. Gershman, S. J. & Uchida, N. Believing in dopamine. Nat. Rev. Neurosci. 20, 703–714 (2019).

10. Reynolds, J. N. J. & Wickens, J. R. Dopamine-dependent plasticity of corticostriatal synapses. Neural Netw. Off. J. Int. Neural Netw. Soc. 15, 507–521 (2002).

11. Richards, B. A. et al. A deep learning framework for neuroscience. Nat. Neurosci. 22, 1761–1770 (2019).

12. Krabbe, S., et al. Adaptive Disinhibitory Gating by VIP Interneurons Permits Associative Learning. http://biorxiv.org/lookup/doi/10.1101/443614 (2018) doi:10.1101/443614.

13. Pi, H.-J. et al. Cortical interneurons that specialize in disinhibitory control. Nature 503, 521–524 (2013).

14. Pinto, L. & Dan, Y. Cell-Type-Specific Activity in Prefrontal Cortex during Goal-Directed Behavior. Neuron 87, 437–450 (2015).

15. Ren, C. et al. Global and subtype-specific modulation of cortical inhibitory neurons regulated by acetylcholine during motor learning. Neuron 110, 2334–2350.e8 (2022).

16. Szadai, Z. et al. Cortex-wide response mode of VIP-expressing inhibitory neurons by reward and punishment. eLife 11, e78815 (2022).

17. Ramamurthy, D. L. et al. VIP interneurons in sensory cortex encode sensory and action signals but not direct reward signals. Curr. Biol. CB 33, 3398–3408.e7 (2023).

18. Gonchar, Y., Wang, Q. & Burkhalter, A. Multiple distinct subtypes of GABAergic neurons in mouse visual cortex identified by triple immunostaining. Front. Neuroanat. 1, 3 (2007).

19. Kim, Y. et al. Brain-wide Maps Reveal Stereotyped Cell-Type-Based Cortical Architecture and Subcortical Sexual Dimorphism. Cell 171, 456–469.e22 (2017).

20. Staiger, J. F., Masanneck, C., Schleicher, A. & Zuschratter, W. Calbindin-containing interneurons are a target for VIP-immunoreactive synapses in rat primary somatosensory cortex. J. Comp. Neurol. 468, 179–189 (2004).

21. Campagnola, L. et al. Local connectivity and synaptic dynamics in mouse and human neocortex. Science 375, eabj5861 (2022).

22. Karnani, M. M. et al. Cooperative Subnetworks of Molecularly Similar Interneurons in Mouse Neocortex. Neuron 90, 86–100 (2016).

23. Lee, S., Kruglikov, I., Huang, Z. J., Fishell, G. & Rudy, B. A disinhibitory circuit mediates motor integration in the somatosensory cortex. Nat. Neurosci. 16, 1662–1670 (2013).

24. Pfeffer, C. K., Xue, M., He, M., Huang, Z. J. & Scanziani, M. Inhibition of inhibition in visual cortex: the logic of connections between molecularly distinct interneurons. Nat. Neurosci. 16, 1068–1076 (2013).

25. de Vries, S. E. J. et al. A large-scale standardized physiological survey reveals functional organization of the mouse visual cortex. Nat. Neurosci. 23, 138–151 (2020).

26. Garrett, M. et al. Experience shapes activity dynamics and stimulus coding of VIP inhibitory cells. eLife 9, e50340 (2020).

27. Ibrahim, L. A. et al. Cross-Modality Sharpening of Visual Cortical Processing through Layer-1-Mediated Inhibition and Disinhibition. Neuron 89, 1031–1045 (2016).

28. Keller, A. J. et al. A Disinhibitory Circuit for Contextual Modulation in Primary Visual Cortex. Neuron 108, 1181–1193.e8 (2020).

29. Khan, A. G. et al. Distinct learning-induced changes in stimulus selectivity and interactions of GABAergic interneuron classes in visual cortex. Nat. Neurosci. 21, 851–859 (2018).

30. Mesik, L., et al. Functional response properties of VIP-expressing inhibitory neurons in mouse visual and auditory cortex. Front. Neural Circuits 09, (2015).

31. Bastos, G. et al. Top-down input modulates visual context processing through an interneuron-specific circuit. Cell Rep. 42, 113133 (2023).

32. Batista-Brito, R. et al. Developmental Dysfunction of VIP Interneurons Impairs Cortical Circuits. Neuron 95, 884–895.e9 (2017).

33. Dipoppa, M. et al. Vision and Locomotion Shape the Interactions between Neuron Types in Mouse Visual Cortex. Neuron 98, 602–615.e8 (2018).

34. Fu, Y. et al. A cortical circuit for gain control by behavioral state. Cell 156, 1139–1152 (2014).

35. Garcia-Junco-Clemente, P. et al. An inhibitory pull-push circuit in frontal cortex. Nat. Neurosci. 20, 389–392 (2017).

36. Kamigaki, T. & Dan, Y. Delay activity of specific prefrontal interneuron subtypes modulates memory-guided behavior. Nat. Neurosci. 20, 854–863 (2017).

37. Luo, X. et al. Synaptic Mechanisms Underlying the Network State-Dependent Recruitment of VIP-Expressing Interneurons in the CA1 Hippocampus. Cereb. Cortex 30, 3667–3685 (2020).

38. Pakan, J. M. et al. Behavioral-state modulation of inhibition is context-dependent and cell type specific in mouse visual cortex. eLife 5, e14985 (2016).

39. Reimer, J. et al. Pupil Fluctuations Track Fast Switching of Cortical States during Quiet Wakefulness. Neuron 84, 355–362 (2014).

40. Williams, L. E. & Holtmaat, A. Higher-Order Thalamocortical Inputs Gate Synaptic Long-Term Potentiation via Disinhibition. Neuron 101, 91–102.e4 (2019).

41. Askew, C. E., Lopez, A. J., Wood, M. A. & Metherate, R. Nicotine excites VIP interneurons to disinhibit pyramidal neurons in auditory cortex. Synapse 73, e22116 (2019).

42. Kepecs, A. & Fishell, G. Interneuron cell types are fit to function. Nature 505, 318–326 (2014).

43. McFarlan, A. R. et al. The plasticitome of cortical interneurons. Nat. Rev. Neurosci. 24, 80–97 (2023).

44. Prönneke, A., Witte, M., Möck, M. & Staiger, J. F. Neuromodulation Leads to a Burst-Tonic Switch in a Subset of VIP Neurons in Mouse Primary Somatosensory (Barrel) Cortex. Cereb. Cortex 30, 488–504 (2020).

45. Rudy, B., Fishell, G., Lee, S. & Hjerling-Leffler, J. Three groups of interneurons account for nearly 100% of neocortical GABAergic neurons. Dev. Neurobiol. 71, 45–61 (2011).

46. Tremblay, R., Lee, S. & Rudy, B. GABAergic Interneurons in the Neocortex: From Cellular Properties to Circuits. Neuron 91, 260–292 (2016).

47. Zeisel, A. et al. Molecular Architecture of the Mouse Nervous System. Cell 174, 999–1014.e22 (2018).

48. Devoto, P. & Flore, G. On the Origin of Cortical Dopamine: Is it a Co-Transmitter in Noradrenergic Neurons? Curr. Neuropharmacol. 4, 115–125 (2006).

49. Lohani, S. et al. Spatiotemporally heterogeneous coordination of cholinergic and neocortical activity. Nat. Neurosci. 25, 1706–1713 (2022).

50. Reimer, J. et al. Pupil fluctuations track rapid changes in adrenergic and cholinergic activity in cortex. Nat. Commun. 7, 13289 (2016).

51. Zaborszky, L. et al. Neurons in the basal forebrain project to the cortex in a complex topographic organization that reflects corticocortical connectivity patterns: an experimental study based on retrograde tracing and 3D reconstruction. Cereb. Cortex N. Y. N 1991 25, 118–137 (2015).

52. Guo, W., Robert, B. & Polley, D. B. The Cholinergic Basal Forebrain Links Auditory Stimuli with Delayed Reinforcement to Support Learning. Neuron 103, 1164–1177.e6 (2019).

53. Ballinger, E. C., Ananth, M., Talmage, D. A. & Role, L. W. Basal Forebrain Cholinergic Circuits and Signaling in Cognition and Cognitive Decline. Neuron 91, 1199–1218 (2016).

54. Eggermann, E., Kremer, Y., Crochet, S. & Petersen, C. C. H. Cholinergic Signals in Mouse Barrel Cortex during Active Whisker Sensing. Cell Rep. 9, 1654–1660 (2014).

55. Lee, S.-H. & Dan, Y. Neuromodulation of brain states. Neuron 76, 209–222 (2012).

56. Liu, C.-H., Coleman, J. E., Davoudi, H., Zhang, K. & Hussain Shuler, M. G. Selective Activation of a Putative Reinforcement Signal Conditions Cued Interval Timing in Primary Visual Cortex. Curr. Biol. 25, 1551–1561 (2015).

57. Crouse, R. B. et al. Acetylcholine is released in the basolateral amygdala in response to predictors of reward and enhances the learning of cue-reward contingency. eLife 9, e57335 (2020).

58. Gritton, H. J. et al. Cortical cholinergic signaling controls the detection of cues. Proc. Natl. Acad. Sci. 113, (2016).

59. Hangya, B., Ranade, S. P., Lorenc, M. & Kepecs, A. Central Cholinergic Neurons Are Rapidly Recruited by Reinforcement Feedback. Cell 162, 1155–1168 (2015).

60. Howe, W. M. et al. Acetylcholine Release in Prefrontal Cortex Promotes Gamma Oscillations and Theta–Gamma Coupling during Cue Detection. J. Neurosci. 37, 3215–3230 (2017).

61. Robert, B. et al. A functional topography within the cholinergic basal forebrain for encoding sensory cues and behavioral reinforcement outcomes. eLife 10, e69514 (2021).

62. Sturgill, J. F. et al. Basal forebrain-derived acetylcholine encodes valence-free reinforcement prediction error. 2020.02.17.953141 Preprint at 10.1101/2020.02.17.953141 (2020).

63. Letzkus, J. J. et al. A disinhibitory microcircuit for associative fear learning in the auditory cortex. Nature 480, 331–335 (2011).

64. Katona, G. et al. Fast two-photon in vivo imaging with three-dimensional random-access scanning in large tissue volumes. Nat. Methods 9, 201–208 (2012).

65. Szalay, G. et al. Fast 3D Imaging of Spine, Dendritic, and Neuronal Assemblies in Behaving Animals. Neuron 92, 723–738 (2016).

66. Jing, M. et al. An optimized acetylcholine sensor for monitoring in vivo cholinergic activity. Nat. Methods 17, 1139–1146 (2020).

67. Berndt, A., Lee, S. Y., Ramakrishnan, C. & Deisseroth, K. Structure-guided transformation of channelrhodopsin into a light-activated chloride channel. Science 344, 420–424 (2014).

68. Allard, S. & Shuler, M. G. H. Cholinergic Reinforcement Signaling Is Impaired by Amyloidosis Prior to Its Synaptic Loss. J. Neurosci. 43, 6988–7005 (2023).

69. Sacramento, J., Costa, R. P., Bengio, Y. & Senn, W. Dendritic cortical microcircuits approximate the backpropagation algorithm. in Proceedings of the 32nd International Conference on Neural Information Processing Systems 8735–8746 (Curran Associates Inc., Red Hook, NY, USA, 2018).

70. Heeger, D. J. Theory of cortical function. Proc. Natl. Acad. Sci. 114, 1773–1782 (2017).

71. Keller, G. B. & Mrsic-Flogel, T. D. Predictive Processing: A Canonical Cortical Computation. Neuron 100, 424–435 (2018).

72. Saxe, A., Nelli, S. & Summerfield, C. If deep learning is the answer, what is the question? Nat. Rev. Neurosci. 22, 55–67 (2021).

73. Kilgard, M. P. & Merzenich, M. M. Cortical Map Reorganization Enabled by Nucleus Basalis Activity. Science 279, 1714–1718 (1998).

74. Zhao, X., Zhang, Y., Yang, L. & Yang, Y. Acetylcholine Release from Basal Forebrain Promotes Cortical Synaptic Plasticity via Somatostatin Interneurons. http://biorxiv.org/lookup/doi/10.1101/2023.07.05.547662 (2023) doi:10.1101/2023.07.05.547662.

75. Phillis, J. W. Acetylcholine release from the cerebral cortex: Its role in cortical arousal. Brain Res. 7, 378–389 (1968).

76. Semba, K. The Cholinergic Basal Forebrain: A Critical Role in Cortical Arousal. in The Basal Forebrain: Anatomy to Function (eds. Napier, T. C., Kalivas, P. W. & Hanin, I.) 197–218 (Springer US, Boston, MA, 1991). doi:10.1007/978-1-4757-0145-6_10.

77. Celada, P., Puig, M. V. & Artigas, F. Serotonin modulation of cortical neurons and networks. Front. Integr. Neurosci. 7, (2013).

78. Hasselmo, M. E. Neuromodulation and cortical function: modeling the physiological basis of behavior. Behav. Brain Res. 67, 1–27 (1995).

79. Hedrick, T. & Waters, J. Acetylcholine excites neocortical pyramidal neurons via nicotinic receptors. J. Neurophysiol. 113, 2195–2209 (2015).

80. Urban-Ciecko, J., Jouhanneau, J.-S., Myal, S. E., Poulet, J. F. A. & Barth, A. L. Precisely Timed Nicotinic Activation Drives SST Inhibition in Neocortical Circuits. Neuron 97, 611–625.e5 (2018).

81. Isaacson, J. S. & Scanziani, M. How inhibition shapes cortical activity. Neuron 72, 231–243 (2011).

82. Adler, A., Zhao, R., Shin, M. E., Yasuda, R. & Gan, W.-B. Somatostatin-Expressing Interneurons Enable and Maintain Learning-Dependent Sequential Activation of Pyramidal Neurons. Neuron 102, 202–216.e7 (2019).

83. Kuchibhotla, K. V. et al. Parallel processing by cortical inhibition enables context-dependent behavior. Nat. Neurosci. 20, 62–71 (2017).

84. Makino, H. & Komiyama, T. Learning enhances the relative impact of top-down processing in the visual cortex. Nat. Neurosci. 18, 1116–1122 (2015).

85. Muñoz, W., Tremblay, R., Levenstein, D. & Rudy, B. Layer-specific modulation of neocortical dendritic inhibition during active wakefulness. Science 355, 954–959 (2017).

86. Xu, H. et al. A Disinhibitory Microcircuit Mediates Conditioned Social Fear in the Prefrontal Cortex. Neuron 102, 668–682.e5 (2019).

87. Yang, J. et al. Functionally distinct NPAS4-expressing somatostatin interneuron ensembles critical for motor skill learning. Neuron 110, 3339–3355.e8 (2022).

88. Yavorska, I. & Wehr, M. Somatostatin-Expressing Inhibitory Interneurons in Cortical Circuits. Front. Neural Circuits 10, 76 (2016).

89. Dudok, B. et al. Alternating sources of perisomatic inhibition during behavior. Neuron 109, 997–1012.e9 (2021).

90. Jiang, X. et al. Principles of connectivity among morphologically defined cell types in adult neocortex. Science 350, aac9462 (2015).

91. Green, J. et al. A cell-type-specific error-correction signal in the posterior parietal cortex. Nature 620, 366–373 (2023).

92. Zeisel, A. et al. Cell types in the mouse cortex and hippocampus revealed by single-cell RNA-seq. Science 347, 1138–1142 (2015).

93. Lourenço, J., Koukouli, F. & Bacci, A. Synaptic inhibition in the neocortex: Orchestration and computation through canonical circuits and variations on the theme. Cortex 132, 258–280 (2020).

94. Bastos, A. M. et al. Canonical Microcircuits for Predictive Coding. Neuron 76, 695–711 (2012).

95. Hertäg, L. & Clopath, C. Prediction-error neurons in circuits with multiple neuron types: Formation, refinement, and functional implications. Proc. Natl. Acad. Sci. U. S. A. 119, e2115699119 (2022).

96. Hertäg, L. & Sprekeler, H. Learning Prediction Error Neurons in a Canonical Interneuron Circuit. http://biorxiv.org/lookup/doi/10.1101/2020.02.27.968776 (2020) doi:10.1101/2020.02.27.968776.

97. Guerguiev, J., Lillicrap, T. P. & Richards, B. A. Towards deep learning with segregated dendrites. eLife 6, e22901 (2017).

98. Lacefield, C. O., Pnevmatikakis, E. A., Paninski, L. & Bruno, R. M. Reinforcement Learning Recruits Somata and Apical Dendrites across Layers of Primary Sensory Cortex. Cell Rep. 26, 2000–2008.e2 (2019).

99. Larkum, M. A cellular mechanism for cortical associations: an organizing principle for the cerebral cortex. Trends Neurosci. 36, 141–151 (2013).

100. Greedy, W., Zhu, H. W., Pemberton, J., Mellor, J. & Ponte Costa, R. Single-phase deep learning in cortico-cortical networks. Adv. Neural Inf. Process. Syst. 35, 24213–24225 (2022).

101. Payeur, A., Guerguiev, J., Zenke, F., Richards, B. A. & Naud, R. Burst-dependent synaptic plasticity can coordinate learning in hierarchical circuits. Nat. Neurosci. 24, 1010–1019 (2021).

102. Urai, A. E. et al. Citric Acid Water as an Alternative to Water Restriction for High-Yield Mouse Behavior. eNeuro 8, (2021).

103. Lerner, T. N. et al. Intact-Brain Analyses Reveal Distinct Information Carried by SNc Dopamine Subcircuits. Cell 162, 635–647 (2015).

104. Wilson, H. R. & Cowan, J. D. Excitatory and inhibitory interactions in localized populations of model neurons. Biophys. J. 12, 1–24 (1972).

105. Vogels, T. P., Sprekeler, H., Zenke, F., Clopath, C. & Gerstner, W. Inhibitory plasticity balances excitation and inhibition in sensory pathways and memory networks. Science 334, 1569–1573 (2011).

106. Urban-Ciecko, J. & Barth, A. L. Somatostatin-expressing neurons in cortical networks. Nat. Rev. Neurosci. 17, 401–409 (2016).

